# Neuropixels reveal structure-function relationships in monkey V1 *in vivo*

**DOI:** 10.1101/2025.05.14.653875

**Authors:** Nicole Carr, Shude Zhu, Xiaomo Chen, Kenji Lee, Alec Perliss, Tirin Moore, Chandramouli Chandrasekaran

## Abstract

The relationship between structural properties of diverse neuronal populations in monkey primary visual cortex (V1) and their *in vivo* functional responses is not fully understood. We combined high-density Neuropixels recordings across cortical layers of macaque V1 with non-linear dimensionality reduction on waveform shape to delineate nine putative cell classes: 4 narrow-spiking (NS), 4 broad-spiking (BS) and 1 tri-phasic (TP). Using targeted analyses of laminar organization, spike amplitude, multichannel waveforms, functional properties, and network connectivity of these cell classes, we demonstrate four aspects of the V1 microcircuit predicted by anatomical studies but never fully demonstrated *in vivo*. First, NS neurons were concentrated in layer 4. Second, a large-amplitude NS cell class in layer 4B showed strong direction selectivity. Third, another layer 4B NS class exhibited robust bursting and orientation selectivity. Finally, cross-correlation analysis revealed functional interactions between cells in different layers. Our results highlight how high-resolution electrophysiology can reveal novel relationships between *in vivo* function of neurons and the underlying circuit.

**Teaser:** High-resolution electrophysiology used with machine learning reveals links between function and the underlying neural circuitry.

## 1. Introduction

The primary visual cortex (V1) of the monkey is the first cerebral cortical locus involved in vision and has been the subject of intense anatomical, *in vitro*, and *in vivo* research over the past 50 years (1–9). A recent single-cell RNA-seq study described 13 transcriptomically separable excitatory cell classes, and 5 inhibitory classes in macaque V1 (10). These different neuron types are located in different laminae and sublaminae (1), possess distinct morphology (11), express a diversity of ion channels (12, 13), and inter-laminar connection patterns (6). Other studies have provided insight into the morphological properties of neurons that connect to other visual areas such as MT and V2 (14–17). In parallel, *in vitro* studies in the cat and monkey have delineated the firing properties of neurons in various layers of cortex (11) and identified bursting neural populations (18, 19). Finally, neurophysiological experiments have identified tuning to diverse visual features (e.g., orientation, color, and direction) in V1 neurons (20–24), and described narrow-spiking bursting cells *in vivo* that are strongly orientation selective (19).

Despite this wealth of research, technical limitations have meant that the majority of these studies examined electrophysiological responses of single neurons *in vivo* to visual stimuli, *in vitro* responses, morphological properties, and laminar organization, separately. *Therefore, the link between functional properties of V1 neurons (e.g., direction selectivity) and their layer and neuronal type-specific properties (e.g., cell size or morphology) is not fully resolved.* We address this question here by capitalizing on three unique capabilities of high-density Neuropixels electrodes: 1) simultaneous recordings from large numbers of neurons, 2) high-resolution recordings across laminae of a brain area, and 3) the ability to record the same neuron on multiple channels. We used these Neuropixels electrodes to record large populations of neurons across all layers of monkey V1, with high spatial resolution, as they respond to visual stimuli (25, 26). We then use our novel WaveMAP machine learning approach (27, 28) to delineate candidate cell types based on the normalized extracellular waveform shape. We then performed a series of targeted analyses of laminar organization, spike amplitude, multichannel spatial features (29), receptive field properties (21), and network connectivity (26) to map the functional responses of these candidate cell types onto underlying microcircuit structure predictions from past research on V1 (13, 14, 19). In this paper, we refer to any *in vivo* electrophysiological responses to stimuli (e.g., tuning, orientation selectivity, direction selectivity, cross-correlations, etc.) as function. Similarly, we refer to any properties intrinsic to the cell (e.g., single– and multichannel waveform shape, laminar distribution, amplitude, inter-spike interval, etc.) as structure.

By leveraging capabilities of high-density Neuropixels recordings with rigorous laminar localization, we discovered four previously unreported links between *in vitro* structure and *in vivo* function: First, anatom-ical studies suggest that in monkey V1, ion channels in the Kv3 family (specifically the Kv3.1b channel), which produce rapid repolarization in narrow-spiking waveforms, are most commonly expressed in layers 4A/B and 4C and are found in both inhibitory and excitatory neurons (mainly in parvalbumin posi-tive, and in a fraction of calbindin positive neurons 12, 13). The studies predict that narrow-spiking neurons should be most common in layer 4 and also more prevalent than the expected 10% fraction of parvalbumin-positive (PV) inhibitory neurons in cortex. Consistent with these predictions, we found that narrow-spiking neurons were most concentrated in layer 4 and occurred more frequently than the expected fraction of PV neurons in cortex, a result aligning with a recent electrophysiological study (19).

Second, anatomical studies show that neurons in V1 projecting to MT and thick stripes of V2, are more likely to be in layer 4B, have large somata, are strongly direction selective, and mostly spiny stellate (12, 14, 15, 17, 23). In accordance with this prediction, we found that neurons with the largest spike amplitudes were found in layer 4B and were strongly direction selective (23, 30, 31). Moreover, the multichannel waveform was more symmetric, consistent with a putative stellate cell morphology (18, 32, 33).

Third, *in vivo* and *in vitro* studies predict the existence of narrow-spiking neurons with strong burst activity (18) with the *in vivo* studies predicting that these neurons are also strongly orientation selective (19). Consistent with this finding, we found a narrow-spiking subpopulation localized to granular layers with 1) inter-spike intervals strongly consistent with bursting, and 2) with the strongest orientation selectivity of all cells recorded.

Fourth and finally, connectivity studies emphasize a feedforward flow of information from layers 4A/B and 4C and 5/6 to layers 2/3 (6). Our cross-correlation analyses revealed that cell populations that were more common in granular and infragranular layers were more likely to lead cell populations in supragranular layers.

In summary, our results demonstrate how large-scale high-resolution laminar electrophysiology enables discovery of relationships between the structural and functional organization of neural responses in V1 *in vivo*. Our approach expands on previous research that suggested a sophisticated microcircuit in monkey V1 and emphasizes that properties such as morphology, laminar organization, and waveform shape are likely correlated to functional properties such as orientation and direction selectivity (31). Finally, these findings suggest that many of the structural details likely result in *in vivo* functional differences, unique to the macaque monkey V1 and should be considered when building new biologically realistic computational models of visual processing in V1 (34). In addition, approaches developed here might facilitate efforts to use Neuropixels to understand relationships between anatomy, biophysical properties, and function in other brain areas.

## 2. Results

Our goal for this study was to use the unique capabilities of Neuropixels to further understand the link between structure and function in the primary visual cortex of monkeys by testing predictions from anatomical studies. To this end, we re-analyzed a recently reported dataset with high-resolution electrophysiological recordings in monkey V1 (25, 26). This dataset is composed of five Neuropixels recordings from the primary visual cortex (V1) of two isoflurane anesthetized macaque monkeys (M1 [sessions 1-3] and M2 [sessions 4-5], Fig. 1A, see *Methods Section: Surgical Details*). The Neuropixels probes were coated with DiI, a dye, to facilitate the histological analysis of each recording location and delineation of layer boundaries. For monkey M1, these analyses suggested that the recordings were performed nearly perpendicular to the lateral opercular surface of V1 and largely restricted to the surface cortex (Fig. S1A, B). For both sessions of monkey M2, the probe penetrated through the surface opercular V1 and white matter, and reached the deep calcarine V1 cortex below (Fig. S1C, D, see *Methods Section: Electrophysiological Recordings*). We then combined these histologically identified boundaries with current source density analysis to assign neurons to layer compartments (see *Methods Section: Scaling Laminar Depths*).

**Figure 1:**
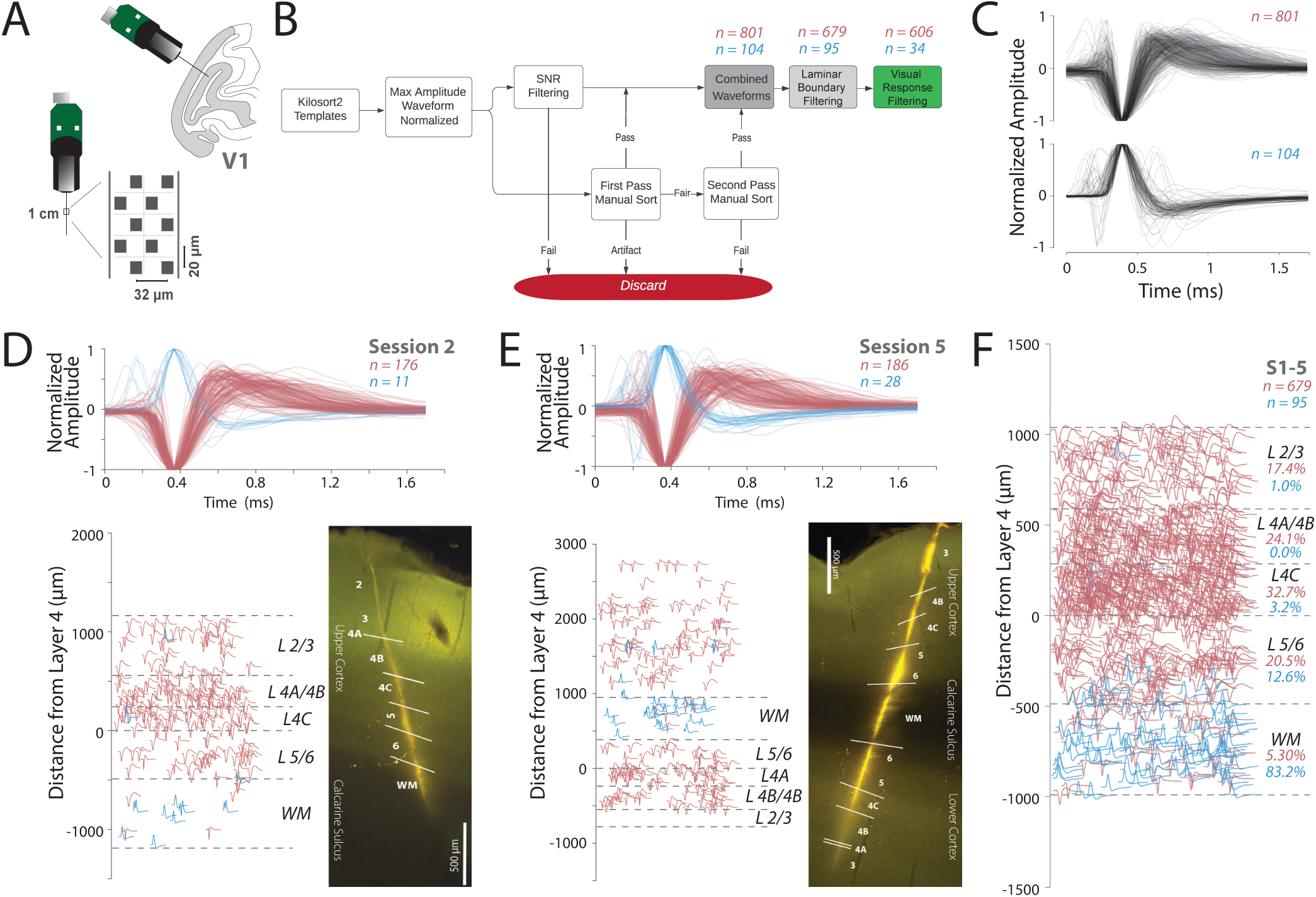
Neuropixels recordings reveal rich neural populations across layers of V1. (**A**) *Upper*, The angle of probe penetrations made into the lateral surface and underlying calcarine sulcus of V1. *Lower*, Diagram of Neuropixels 1.0 probe base and shank. Right diagram shows the layout of electrode contacts for a section of the recording shank. (**B**) Semiautomatic quality control process applied to single unit templates extracted from Kilosort2 analysis to identify single neurons. This process includes the following steps: 1) Maximum amplitude channel of extracellular waveform template averaged over all identified spikes for each neuron and then normalized between –1 and 1. 2) Each single unit sorted by SNR thresholding is manually curated. We used two independent observers for manual curation of the waveform templates with an inter-rater reliability of 0.72. 3) Waveforms that passed SNR filtering and manual curation form the final dataset (*pink*: negative spiking units, *blue*: positive spiking units). (**C**) *Upper*, 801 negative spiking extracellular waveforms after quality control. *Lower*, 104 positive spiking extracellular waveforms. (**D**) Example Session 2, Monkey 1 (M1), *Upper*, 187 waveforms across all layers of cortex in primary visual cortex (*pink*: negative spiking units, *blue*: positive spiking units). *Lower Left*, laminar distribution with marked layer boundaries, determined by current source density and histology. *Lower Right*, histology image of probe track. (**E**) Example Session 5, Monkey 2 (M2), *Upper*, 214 waveforms across all layers of cortex in primary visual cortex (*pink*: negative spiking units, *blue*: positive spiking units). *Lower Left*, laminar distribution with marked layer boundaries, determined by current source density and histology. The layers here are flipped because the session 5 recording was taken from the deeper cortex. *Lower Right*, histology image of probe track through calcarine sulcus into cortex below. (**F**) Pooled waveforms within laminar boundaries from M1 and M2, sessions 1-5, *Upper*, 774 waveforms across all layers of cortex in primary visual cortex with scaled depths (*pink*: 679 negative spiking units, *blue*: 95 positive spiking units).

During each recording, neurons in V1 were stimulated with drifting Gabor gratings in 36 directions, 4 different spatial frequencies, and either monocular or monocular and binocular conditions (see *Methods Section: Visual stimulation*). While these stimuli were displayed to the animal, we recorded broadband extracellular neural activity from V1 and only used the optimal eye conditions for the analysis of functional properties. We used Kilosort2 to spike sort the broadband extracellular data and extract templates representing the waveform of the extracellular action potential from each unit (35).

After spike sorting these five recording sessions, we obtained a total of 2,529 templates. We then used a rigorous and conservative waveform curation process, by combining signal-to-noise ratio (SNR) filtering and manual inspection of the maximum amplitude action potential waveform. We only considered spikes that were measured on multiple electrode channels. We also manually identified and removed artifacts, noisy units, and units outside the laminar boundaries to identify a conservative single neuron population (Fig. 1B, see *Methods Section: Spike Sorting and Data Curation*).

After this semi-automated curation process, we were left with 905 single neurons over the five sessions (Fig. S1E). We then separated neurons into “positive spiking” waveforms, where the maximum (hyper-polarization) peak came before the minimum (depolarization) trough, and “negative spiking” waveforms, where the minimum trough came before the maximum peak. The positive spiking neurons were aligned to the peak, and the negative spiking neurons to the trough. In the case of tri-phasic waveforms, where two major peaks were detected on either side of the trough, the waveforms were aligned to the trough and second peak, and were classified as “negative spiking” waveforms.

We identified 801 negative spiking neurons, and 104 positive spiking neurons from all recording sessions (Fig. 1C, upper and lower panels respectively, Fig. S1E). In sessions 4 and 5 of monkey M2, we excluded the neurons from surface V1, because the visual stimulation was performed based on the RF locations of the calcarine sulcus V1 (Fig. S1C-D,G). Since we were comparing the waveform to the functional properties, we only selected neurons that were responsive to visual stimuli. For instance, this would mean that in Fig. 1E, waveforms from +1000 *µ*m to 3000 *µ*m were excluded from the analysis. Of the negative spiking units, 606 were both responsive to visual stimuli and within the layer boundaries from layers 2/3 to white matter (Fig. S1F-G). These negative spiking units were used as the primary dataset for linking structure and function. The positive spiking units had very few visually responsive units, and thus were not included in functional analyses; however, they did show consistent laminar organization as discussed in a separate section (see *Results Section: 2.7*).

Overall, the negative waveforms were more common in gray matter (layers 1-6, pink, *χ*^2^ (1, 679) = 542.635, p *<* 0.0001, Fig. 1F, Fig. S1F-G), whereas the positive waveforms were more common in white matter (blue, *χ*^2^ (1, 95) = 41.779, p *<* 0.0001, Fig. 1F). For example, in session S2 from monkey 1, the recordings were largely restricted to the opercular surface of V1, and the positive waveforms were largely found in the white matter (blue, Fig. 1D). Additionally, even with sessions (4–5) from monkey 2, in which the recordings spanned both the opercular cortex, the white matter, and the cortex below the calcarine sulcus, positive waveforms were more common in the white matter (Fig. 1E).

Fig. 1F summarizes this pattern from all five sessions in a normalized space with layer boundaries. To preserve the depth positions of neurons within their original layer boundaries (see *Methods Section: Scaling Laminar Depths*), we first calculated the average layer boundaries of the five sessions. We scaled the depths of each neuron to fit within the average layer boundaries so that all five sessions could be analyzed together. Then, for each neuron in each session, we identified the layer in which the neuron was present. Orientation tuning curves for all the visually responsive neurons (Fig. S1H) were similar suggesting that our recordings were largely perpendicular to the cortical surface (25, 26).

We examined the overall visual response properties (see *Methods Section: Receptive Field Properties*) of this V1 dataset, including tuning and latencies, and confirmed that they robustly replicated previous results from V1 (21). First, we validated that layer 4C neurons responded earlier than all other layers (36, (Fig. S3A)). In addition, we found both simple and complex cells in V1 and these simple cells were more common in layers 4A/B and 4C, similar to previous reports in V1 (21, (Fig. S3B, D)). These findings reassured us that the dataset was consistent with many of the previous findings in V1. In the next sections, we take advantage of these rich high-resolution laminar recordings and go beyond these established findings. We delineate candidate cell types from the shape of the waveforms recorded and analyze various properties to derive deeper links between structure and function.

### 2.1. WaveMAP identifies broad and narrow-waveform clusters from negative spiking units

We analyzed the negative and positive spiking waveforms separately using our previously published WaveMAP approach (27, 28) to delineate putative cell types that could be subjected to further analysis (see *Methods Section: WaveMAP Analysis*). In this approach, we used uniform manifold approximation and projection (UMAP) to first create a high-dimensional graph, followed by Louvain clustering to de-lineate putative cell types, and finally projected into 2D space for visualization of the high-dimensional structure. This non-linear dimensionality reduction technique outperforms the traditional approaches that rely on a small number of classical features, such as trough-to-peak duration, spike width, or repolarization slope, and has been used by several others to identify candidate cell types (27, 38, 39).

Although waveform shape is known to depend on the location of the probe along the morphological structure of the neuron, most biphasic extracellular action potentials (EAPs) recorded from electrode channels are likely to be somatic, while tri-phasic waveforms are more associated with dentritic return currents or axons (39–42). In this high-density Neuropixels dataset, where each neuron was captured across multiple recording channels, we selected the waveform with maximum amplitude for analysis, thus centering our analysis on EAPs most likely to be somatic. Then as a pre-processing step for input into WaveMAP, we normalized the waveforms, which emphasizes aspects of the waveform including transitions in trough-peak width and repolarization width, that emerge from activity-dependent ion channel kinetics associated with different cell types. We found that the classical features of width, trough-to-peak distributions, and amplitude were captured by WaveMAP, but were largely unimodal (Fig. S2C, E). While not easily separated by traditional approaches, this Neuropixels dataset was more logically delineated using the unsupervised clustering in high-dimensional space.

Fig. 2A shows the result of the WaveMAP analysis on 801 negative waveforms with a resolution parameter of 1.0 for Louvain clustering. WaveMAP, like all machine learning algorithms, benefits from large and high-quality datasets. To improve the results of the WaveMAP clustering, we included all 801 negative spiking waveforms, then chose the 606 neurons with visual responsiveness from this population for further analysis in linking structure and function. We found 9 clusters captured the heterogeneity of waveform shapes in this dataset. The cluster shapes were either narrow and tri-phasic (TP-1, green, Fig. 2A), narrow and biphasic with a sharp after hyperpolarization phase (Clusters NS1-4, cooler colors, Fig. 2A) or broad and biphasic with a slower after hyperpolarization phase (Clusters BS1-4, warmer colors, Fig. 2A). We quantified the fraction of neurons in each cluster and each general category (see inset). Most of the neurons had broad-spiking waveforms (54%). A smaller percentage of neurons had narrow, tri-phasic (14%) waveforms, and narrow-spiking waveforms made up the remaining sample (32%).

**Figure 2:**
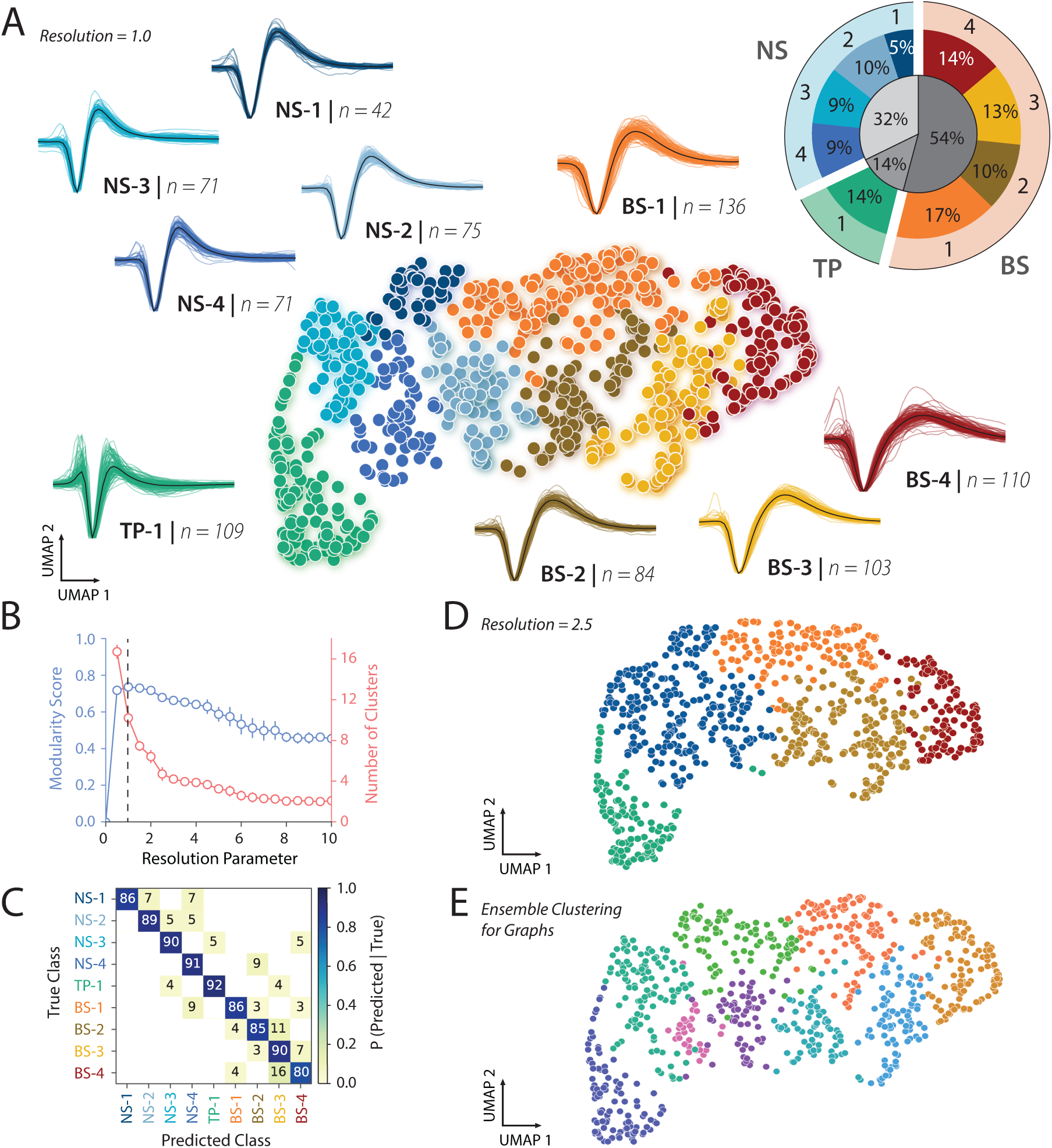
UMAP and Louvain Clustering on 801 negative spiking waveforms reveals 9 candidate cell types. (**A**) Scatter plot of waveforms in UMAP space colored by Louvain cluster membership. WaveMAP parameters were set as the following: N_neighbors = 20; MIN_DIST = 0.2; RESOLUTION = 1.0. Cooler colors denote narrow-spiking clusters, and warmer colors denote broad-spiking clusters. Adjacent to each numbered cluster is shown all member waveforms and the average waveform shape (in black). Each window is 1.8 ms in duration. *Inset*, Population percentages by cluster. (**B**) The Louvain clustering resolution parameter plotted against the resulting modularity score (in blue, on left axis). On the same plot, the Louvain clustering resolution versus the number of clusters (in red, on right axis). This was averaged over 25 runs for WaveMAP using 25 random samples and seeds of 80% of the full dataset at each resolution parameter from 0 to 10 in 0.5 unit increments (a subset of the data was used to obtain error bars). Each data point is the mean ± standard deviation. The vertical dashed line indicates the maximization of modularity (27). (**C**) The confusion matrix of a gradient boosted decision tree classifier with five-fold cross-validation. The main diagonal shows accuracy of waveform classification for each cluster, and off diagonals show misclassification percentages (27). The average accuracy across all clusters was 88%. (**D**) Scatter plot of waveforms in UMAP space colored by Louvain cluster membership with a larger resolution parameter. Conventions as in (A). WaveMAP parameters were set as the following: N_neighbors = 20; MIN_DIST = 0.2; RESOLUTION = 2.5. (**E**) An alternative version of the Louvain clustering algorithm (ensemble clustering for graphs (37), requires setting no resolution parameter, and produces 9 clusters similar to our clustering at RESOLUTION = 1.0.

We chose the resolution parameter for the Louvain clustering using the following approach. We calculated the modularity score which is the ratio of graph connectivity within a cluster to the connections from outside into a cluster (Fig. 2B). Maximizing modularity optimized the community “connectedness” (27). We found that a resolution parameter of 1.0 maximized the modularity score (Fig. 2B, see *Methods Section: WaveMAP Analysis*).

The resolution parameter tuning is important because if it is too fine, this leads to over-clustering and producing clusters very similar to each other. To assess whether this was the case, we performed a five-fold cross-validation analysis to understand how separable these clusters were from one another. The confusion matrix from the classification analysis suggested 1) that the clusters were actually quite well separated with a mean accuracy of 88%, and 2) with minimal misclassification (Fig. 2C). This suggests that our choice of the resolution parameter was a good balance between maximizing the modularity score and not over-clustering.

Consistent with these results, when we used an alternative Ensemble Clustering for Graphs (ECG) ap-proach that does not require setting a fixed resolution parameter, we again found that the data are well explained by 9 clusters (Fig. 2E). Finally, the Louvain clustering method used in WaveMAP is also a hi-erarchical method. Decreasing the resolution parameter does not lead to an entirely new clustering result but rather some clusters are merged together while other clusters are kept intact. In our case, clusters NS1-4 were merged into one narrow-spiking waveform cluster, clusters BS-2 and BS-3 were merged into one broad-spiking waveform cluster, and clusters BS-1, BS-4, and TP-1 remained independent and stable (Fig. 2D). This is important to note, considering how resolution parameters 0.5 through 2.0 produce similar modularity scores, yet the number of resulting clusters differ drastically. Therefore, we also define general categories that capture more global characteristics than the local features of the 9 clusters, with NS1-4 as the narrow-spiking group, BS1-4 as the broad-spiking group, and TP-1 as the tri-phasic group. This helped validate our choice of resolution parameter and the resulting nine candidate cell classes.

As a final check, we also tested whether the UMAP dimensions and identified clusters were biologically meaningful. If the clusters are biologically meaningful, then the UMAP dimensions/clusters and depth combined should explain more variance in the functional properties than just the depth at which a neuron is found. Consistent with this hypothesis, a linear multiple regression analysis revealed that depth and UMAP clusters explained more variance (as measured by adjusted *R*^2^) in functional properties compared to just depth alone (Fig. S2H). Similarly, depth and candidate cell type explained more variance than depth alone. Finally, these effects were not a trivial artifact of just signal-to-noise ratio as it only explained a small fraction of the variance and far less than the variance explained by UMAP and depth/candidate cell type and depth.

Together, the clustering analyses suggest that nine clusters balance the diversity of candidate cell types while minimizing overclustering of this dataset. In the next sections, we analyze the properties of these various clusters and reveal links between structure and function.

### 2.2. Narrow-spiking waveforms are more likely in granular layers, and more common than PV neurons

Anatomical studies suggest that potassium channels (e.g., in the Kv3 family) that lead to rapid repolarization and narrow-spiking waveforms are more prevalent in layers 4A, 4B, and 4C than in other layers of V1 (13, 43), and although commonly associated with fast-spiking PV cells, are found in both excitatory and inhibitory neurons (12, 44, 45). These Kv3 positive, excitatory neurons are largely restricted to layers 4B and 4C (13, 43). The direct functional prediction from these anatomical studies is that 1) narrow-spiking neurons should be more common in layers 4A/B and 4C compared to other layers, 2) they should be more common than PV neurons, and 3) there should be a subpopulation largely restricted to layers 4A/B and 4C with narrow-spiking waveforms. To test these predictions, we performed three analyses described below.

First, we assessed the laminar distribution of NS1-4 clusters identified from our recordings (Fig. 3), and found that all narrow-spiking classes are more common in layers 4C and 4A/B compared to other cortical layers (Fig. 3, *left panel, density distributions*, Fig. S4A, B, Table S1). Under the null hypothesis, the number of narrow-spiking neurons in all layers should be evenly distributed. In contrast, the alternative hypothesis suggests that narrow-spiking neurons should be more common in layers 4A/B and 4C than in layers 2/3, and layers 5/6. Consistent with the alternative hypothesis, 77.5% ± 2.97% of NS neurons were found in layers 4A/B and 4C, whereas only 12.5% were found in L2/3 and 10% were found in layers 5/6 (*χ*^2^ (2, 199) = 173.97, p *<* 0.0001, (Fig. S4B)). Additionally, the mean distance from layer 4 for all narrow-spiking neurons was 304 *µ*m (95 CI: 266.28, 342.04*µ*m) and was well within the boundaries of layers 4A/B and 4C (0*µ*m – 588.8*µ*m). These two analyses provide evidence that the narrow-spiking neurons are common in layers 4A/B and 4C compared to other layers. This *in vivo* finding, enabled by the high yield of neurons and increased spatial resolution of the high-density electrodes used in this study, supports a prediction made by anatomical studies (13), and thus an example of the link between structure and function.

**Figure 3:**
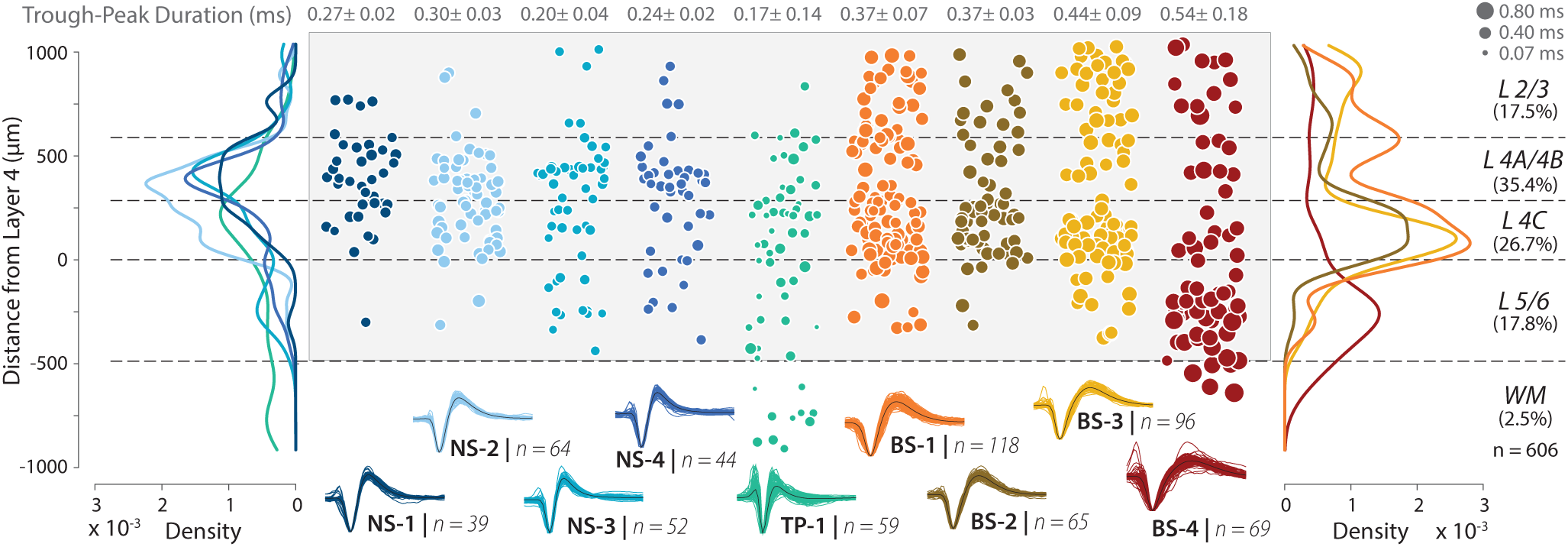
Narrow-spiking waveforms are more common in layers 4A/B and 4C. *Center*, Panel (shaded in light gray) shows a scatter plot of laminar location for neurons in each of the clusters. The mean and the standard deviation of the width of the waveforms are indicated at the top of each plot. Dashed lines depict layer boundaries identified from CSD and immunohistochemistry. Size of the markers indicates trough-to-peak duration. *Left* and *Right* panels show the density distributions for the cell populations. Percent of units in each layer shown in parentheses (shown on the right).

Second, we compared the percentage of narrow-spiking neurons, composed of NS1-4, in the final dataset (606 units) with the percentage of PV neurons expected from anatomical studies of V1. Our estimate of narrow-spiking neurons was 32% ± 1.6% (mean ± sem), which was significantly above the fraction of PV neurons reported in the literature (5-10%, confidence intervals do not overlap) (13, 45). Together, these results are consistent with our predictions from anatomy that narrow-spiking neurons should be more common in layers 4A/B and 4C and likely observed in both excitatory and inhibitory neurons.

Third, we assessed the laminar distribution of each of the NS1-4 subclasses to examine if a subpopulation was largely restricted to layers 4A/B and 4C (Fig. S4C, D), which may account for the excitatory Kv3 positive neurons previously found in anatomical studies (13). Consistent with the structural prediction, 86.4% (*χ*^2^ (2, 103) = 131.33, p *<* 0.0001, Fig. S4D) of NS-1 and NS-2 subclasses was only observed in layers 4A/B and 4C and far less prevalent in layers 2/3 (9.7%) and 5/6 (3.9%). In contrast, although NS-3 and NS-4 were strongly concentrated in layers 4A/B and 4C (67.7%, *χ*^2^ (2, 96) = 52.94, p *<* 0.0001, Fig. S4D), they were also found in layers 2/3 (10.47%) and layers 5/6 (21.9%). Together, the laminar distribution of narrow-spiking neurons is the first previously unreported link between anatomically predicted cell populations and WaveMAP identified candidate cell types in V1.

### 2.3. The largest amplitude neurons in V1 are located in layer 4B and show strong direction selectivity

Anatomical studies of V1 suggest that a subpopulation of stellate neurons in layer 4B with dense dendritic arborization and some of the largest soma sizes project to MT (14, 46), and thick stripes of V2 (17). In parallel, another study (23) used antidromic stimulation to identify neurons that project from V1 to MT and found these neurons to be strongly selective to the drift direction of Gabor gratings. Together, these studies predict a subpopulation of neurons strongly localized to layer 4B with large amplitudes and strong direction selectivity.

We tested this prediction by examining the spike amplitude of each of the cell classes and their direction selectivity. We used spike amplitude as an estimate of cell size based on observations that spike amplitude 1) is related inversely to input resistance with larger neurons having smaller resistance and 2) positively correlated to the size of the soma and the dendritic arbor (47). The null hypothesis is that there is no relationship between spike amplitude and direction selectivity. Alternatively, if there is a population of putative MT (and likely V2) projecting neurons with large soma and extensive dendritic arbors, we should expect that this cluster should 1) have the largest spike amplitudes, 2) be more likely in 4B, and 3) be selective for direction. Note that there is no guarantee that there is a uniform distance from each unit to the recording probe. However, such an effect will minimize relationships between spike amplitude and cluster and lead to a more uniform distribution of cell sizes as a function of cluster, and thus make it harder to reject the null hypothesis.

We tested this prediction using the following analysis. Fig. 4A shows a scatter plot of direction selectivity as a function of candidate cell type and layer. Amplitude varied widely as a function of cell type and layer. We found a modest positive correlation between amplitude and direction selectivity while controlling for the effects of SNR using partial correlation (partial Pearson correlation controlling for SNR, r = 0.2057, p < 0.001). Moreover, consistent with our anatomical predictions, visualization suggested that the largest amplitudes were more common in layer 4B and that these neurons were also strongly direction selective (Fig. 4A, see gray circle).

**Figure 4:**
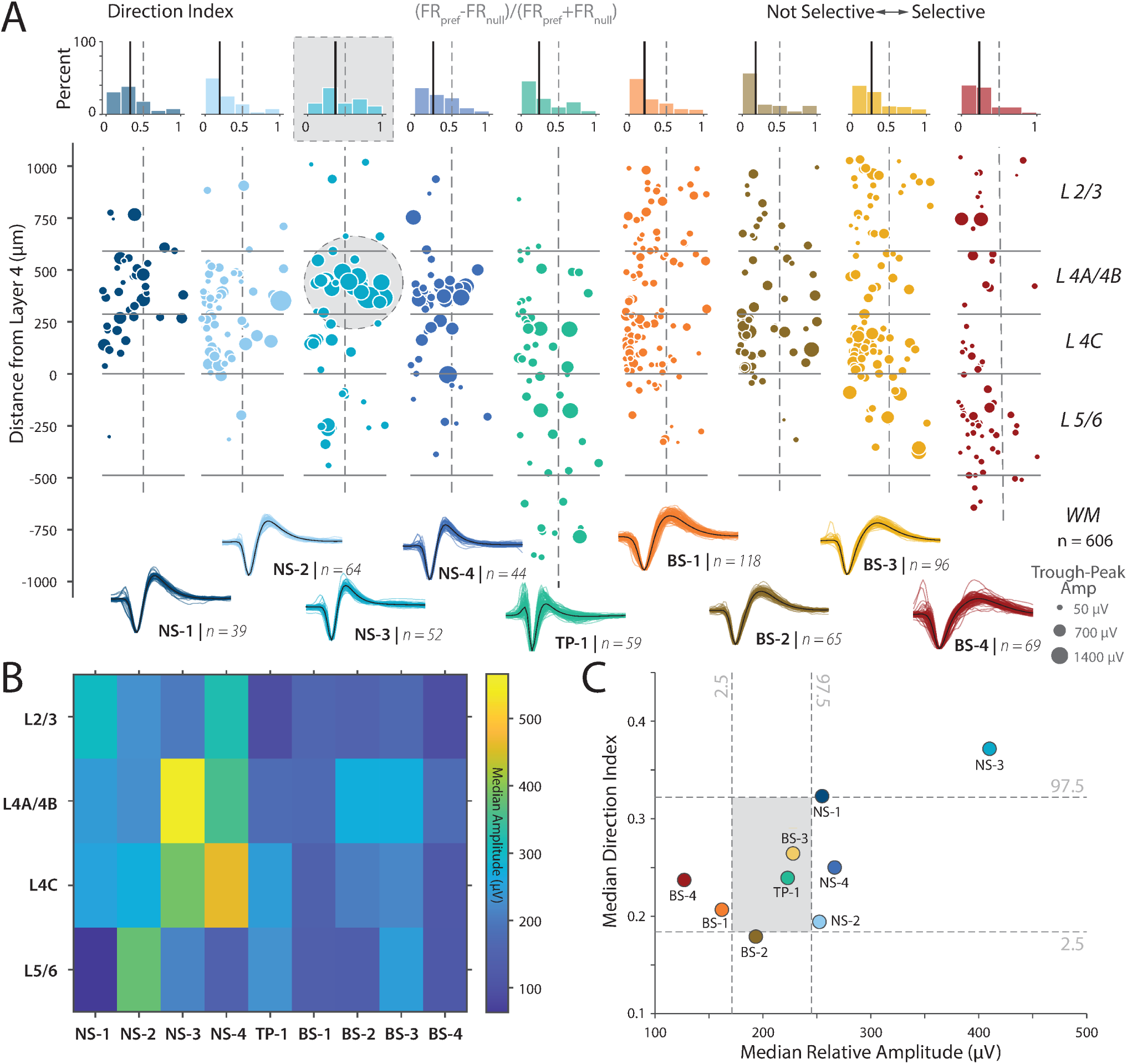
The NS-3 cluster shows strong direction selectivity and high amplitude. (**A**) *Center*, panel shows a scatter plot of direction selectivity (x-axis) vs. depth (y-axis) for visually responsive neurons in each of the clusters. The points are each colored by the cluster. The marker size for neurons within each cluster is scaled by the amplitude of each unit (*µ*V). Solid lines depict layer boundaries identified from CSD and immunohistochemistry. *Top*, panel above the scatter plots shows histograms of the direction selectivity index. The bold line indicates the median direction index for each cluster. The vertical dashed center lines show the boundary between not selective (left of center) and selective (right of center). (**B**) Heatmap of relative trough to peak amplitudes as a function of cluster and layer. (**C**) Plot of median direction index and cluster relative amplitude (*µ*V) for each of the clusters along with the 2.5 and 97.5 percentile for the shuffled distributions estimated from 500 shuffles. Only the NS-3 class has median amplitude and direction index significantly different from the shuffle, whereas only the amplitudes of NS-4, BS-1 and BS-2 are significantly different from the shuffle.

We quantified these qualitative patterns by measuring the median amplitude of the waveforms as a function of cluster and layer (see Fig. 4B). Consistent with both the qualitative picture obtained from (Fig. 4A) and our prediction, the largest amplitude neurons were found in layer 4B (yellow box in colormap). If our hypothesis that these large neurons in layer 4B project to MT is correct, then they should be strongly direction selective (23). We plotted the median amplitude and direction selectivity per cluster (Fig. 4C). Consistent with our prediction, we found that the NS-3 cluster, largely localized to layer 4B, had the largest amplitude and the strongest direction selectivity. We found that the median direction selectivity and the amplitude for the NS-3 class were significantly different from the shuffled distribution (Shuffle test, p < 0.004 for both comparisons for NS-3 but not NS-4 or other classes).

Examining the broad-spiking neurons revealed that clusters BS-4 and BS-1 have the smallest spike amplitudes and were significantly different from the shuffled distribution (Fig. 4C). Cluster BS-4 is largely localized to layers 5/6, whereas cluster BS-1 is most common in the 4C layers (Fig. S4D). These findings are also consistent with the following anatomical observations: First, electron microscopy studies suggest that in layer 5 of V1, most pyramidal neurons are small (48). Second, many studies demonstrate that layers in 4C are composed primarily of stellate neurons which range from small to large stellate cells, but overall the average soma size is smaller in the 4C layers compared to layers 2/3 and layer 4B (48). Cluster BS-2 and BS-3 are both slightly larger in amplitude and more concentrated in layer 4C and layers 2/3. However, cluster BS-2 has significantly less direction selectivity than cluster BS-3. This supports the result that the broad-spiking group is putatively excitatory, and can be separated into functional subtypes by layer and waveform cluster.

While these results are contingent on links between voltage amplitude and neuron size (31), these results are remarkably consistent with predictions from anatomical studies and is evidence for a functional population of neurons in layer 4B with large spike amplitudes and strong direction selectivity.

### 2.4. A narrow-spiking cluster, NS-1, is localized to layer 4B and has “bursting”-like properties

*In vivo* and *in vitro* studies of V1 predict neural populations with bursting activity in V1 (18, 19, 49, 50). One *in vitro* study in the cat predicts that narrow-spiking neurons with strong bursting activity should be found in layers 2-4 (18), and broad-spiking neurons with bursting activity should be observed in all layers. Furthermore, the *in vivo* studies suggest that these bursty, narrow-spiking neurons should be strongly tuned to orientation (19). One signature of “bursting” is peaky inter-spike interval (ISI) distributions. Collectively, these studies predict the existence of narrow-spiking clusters in layers 2-4 with strongly peaked ISI distributions and strong orientation selectivity.

Fig. 5A shows the ISI probability distribution for one narrow-spiking and two broad-spiking example neurons in our dataset. Two of these neurons had peaked ISI distributions, whereas the third had a wide distribution with no clear peak. We calculated the normalized ISI distributions for visually responsive neurons, sorted by the location of the ISI peak for each of the clusters (Fig. 5B, see *Methods Section: Inter-spike Interval*). This plot suggests that the neural population in V1 is heterogeneous and includes neurons with peaked ISI distributions, and those with broad ISI distributions. However, some of the narrow-spiking clusters are more likely to have peaky ISI distributions compared to some of the broader-spiking clusters (e.g., NS-1, compared to BS-3).

**Figure 5:**
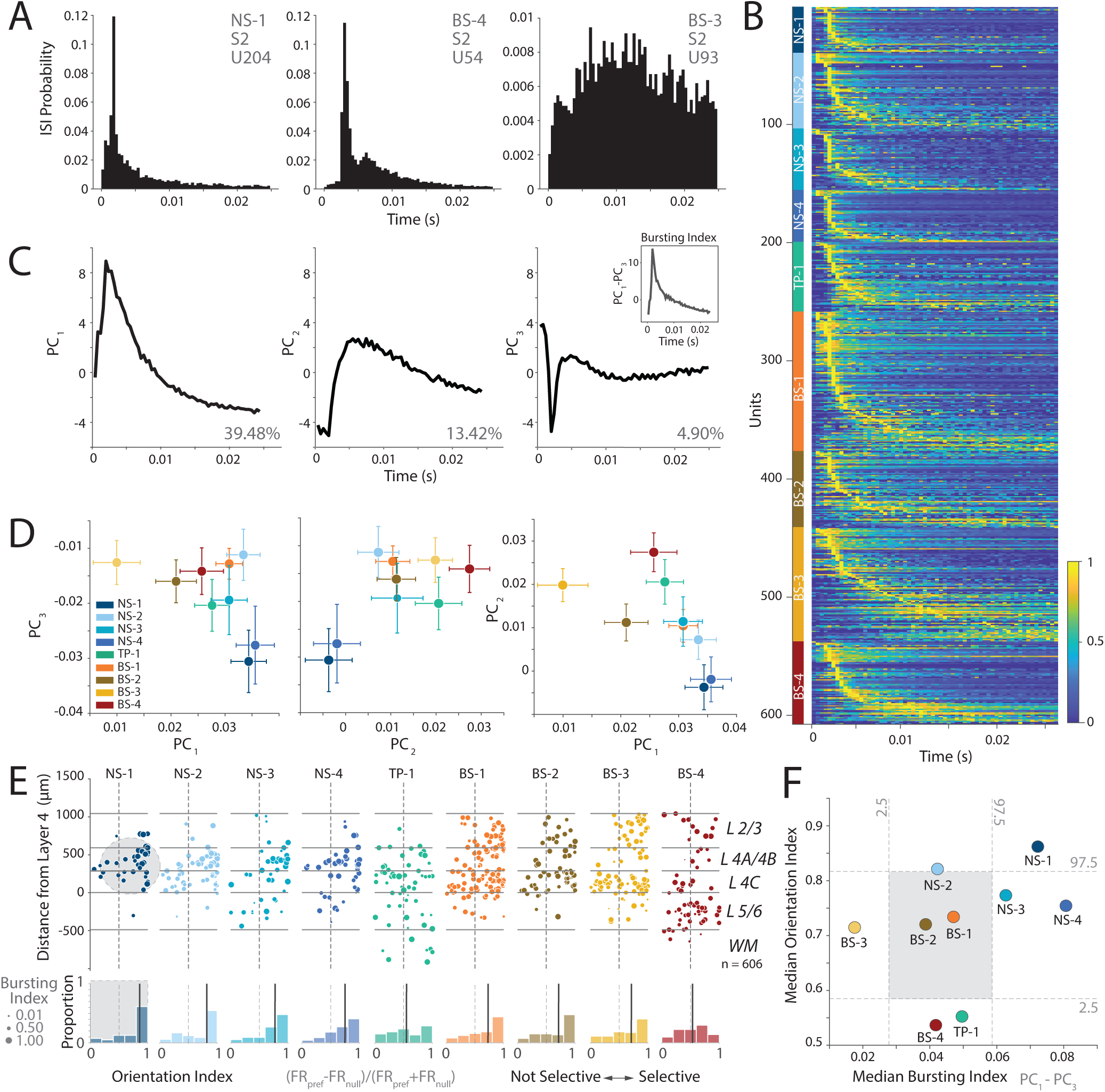
Narrow-spiking, bursty population localized to V1 layers 4A/B and 4C show high orientation selectivity. (**A**) Inter-spike interval histogram normalized to relative probability of three example units from cluster NS-1, BS-4, and BS-3. (**B**) Heatmap of normalized ISI distributions of all visually responsive units, sorted by ISI peak, and by cluster. (**C**) First three PCs of the ISI distributions. *Inset*, The difference between *PC*_1_ and *PC*_3_ emphasize the “peaky” shape of an ISI distribution consistent with bursting. (**D**) *Left*, Scatter plot of mean cluster values for *PC*_1_ and *PC*_3_. *Middle*, Scatter plot of mean cluster values for *PC*_2_ and *PC*_3_. *Right*, Scatter plot of mean cluster values for *PC*_1_ and *PC*_2_. Errorbars indicate SEM. (**E**) *Top*, Panel shows a scatter plot of orientation selectivity (x-axis) vs. laminar location (y-axis) for neurons in each of the clusters. The marker size for neurons within each cluster is scaled by the Bursting Index showing the most “peaky” ISI distributions. Solid lines depict layer boundaries. *Bottom*, Panel below the scatter plots shows histograms of the orientation selectivity index. The bold line indicates the median orientation index for each cluster. The vertical dashed center lines show the boundary between not selective (left of center) and selective (right of center). (**F**) Scatter plot of median orientation index and median bursting index (difference *PC*_1_ and *PC*_3_) per cluster for each of the clusters along with the 2.5 and 97.5 percentile for the shuffled distributions estimated from 500 shuffles. Cluster NS-1 has significantly different orientation selectivity and bursting from the shuffled distributions.

To better understand this heterogeneity, we performed PCA on the normalized ISI distributions. We first examined the components of the ISI distribution and found that *PC*_3_ had a strong dip for short ISIs. Thus, ISI distributions that are more bursting are likely to have a strong negative loading on this principal component.

In contrast, both *PC*_1_ and *PC*_2_ were more similar to ISI distributions expected from regular spiking activity with refractory periods (Fig. 5C). If the *in vitro* prediction is accurate (18), suggesting the presence of a population of narrow-spiking, bursting neurons with high orientation selectivity in V1, then we expect some narrow-spiking clusters with strong negative loadings on *PC*_3_ and positive loadings on *PC*_1_. Consistent with this prediction, both NS-1 and NS-4 clusters had negative loadings on *PC*_3_ and positive loadings on *PC*_1_ suggesting that these units were more likely to be bursting (Fig. 5D). In contrast, BS-3 had minimal to no loading on *PC*_3_ suggesting that these neurons were more likely to be regular spiking. We calculated a bursting index, defined as the difference in loadings between *PC*_1_ and *PC*_3_ for each of the clusters to capture this trend (see *Methods Section: Inter-spike Interval*).

We examined whether there are any relationships between orientation selectivity and the bursting index of each of these clusters, and tested the hypothesis that the bursting narrow-spiking neurons are strongly orientation selective (19). Fig. 5E shows a scatter plot of the orientation selectivity of neurons for each of the clusters, with the dot size a function of the burst index for each neuron. Overall, there was a modest relationship between the bursting index and orientation selectivity, while controlling for the effects of SNR (partial correlation controlling for SNR (Pearson’s r = 0.1250, p < 0.01). Moreover, some clusters were more likely to be strongly bursting and orientation selective. NS-1 had a strongly peaked distribution for orientation selectivity and higher burst indices consistent with the predictions from literature (19). In contrast, NS-4 had broader distribution of orientation selectivity while being likely to have a ISI distribution consistent with bursting. Similarly, NS-2 showed some orientation selectivity but was less likely to show peaked ISIs that are consistent with bursting. Note that the mean firing rate of all of these cells was largely similar (~20 spikes/s, Fig. S3A, C), suggesting that these ISI distributions are the result of rapid “burst” like patterns of spiking in response to the sensory stimulus, and not just a trivial effect of the firing rate.

Finally, we tested if these differences were meaningful by plotting the median orientation index against the median bursting index. The bursting index for NS-1 was significantly different from the shuffled distribution (Shuffle test, p < 0.02, Fig. 5F) indicating that the ISI distributions for these neurons were more peaky and therefore suggesting a greater propensity for bursting. In contrast, BS-3 had the lowest bursting index suggesting that this cluster was more likely to be regular spiking on average (Shuffle test, p < 0.02).

Collectively, these results suggest that bursting activity is observed in all clusters. However, a subpopula-tion of narrow-spiking neurons in V1 (NS-1), on average are 1) more likely to have peaky ISI distributions consistent with bursting, 2) localized to layers 2-4, and 3) orientation selective. These cell populations are an *in vivo* discovery of cell populations predicted by ex-vivo electrophysiological studies of cat V1 (18), and provide laminar localization of bursting narrow-spiking neurons in V1 (19).

### 2.5. NS-1 and NS-3 clusters have distinct multichannel waveforms consistent with candidate pyramidal and stellate neuron morphology

Thus far we have focused on the single channel waveform and attempted to link it to functional properties. However, the high-resolution recordings from Neuropixels provide the unique ability to localize neurons across multiple electrodes and thus can measure a multichannel waveform for a neuron (see *Methods Section: Multichannel Profiles*). Modeling and experimental studies suggest that the multi-channel waveform is at least in part influenced by the morphology of the cell (29, 33). In this section, we examine whether the multichannel waveforms of these neuronal clusters are consistent with anatomical descriptions of the morphology of various cell types reported in V1 (29, 33).

In Fig. 5, we identified that the NS-1 cluster was strongly bursting and orientation selective. These narrow-spiking, bursting cells are thought to be excitatory and pyramidal in nature (18). A recent Neuropixels study of mouse V1 (29) suggested that pyramidal neurons in V1 show evidence of strong uni-directional propagation of action potentials towards the dendrites whereas inhibitory and spiny stellate neurons are more likely to have symmetric waveforms. Thus, if the NS-1 cluster is an excitatory pyramidal cell type, it should show unidirectional propagation of action potentials. Similarly, the NS-3 cluster had large amplitude and high direction selectivity, which were remarkably consistent with anatomical findings about neurons that project from V1 to MT (14). An additional prediction from this study is that the neurons that project from V1 to MT are likely to be stellate in nature (14). Again, modeling studies (33) suggest that non-pyramidal neurons lacking large apical dendrites and instead possessing dense arborization around the soma (e.g., spiny stellate cells in layers 4A, 4B, and 4C) are more likely to have symmetric multichannel waveforms.

We first tested the prediction that NS-1 is an excitatory pyramidal neuron and will show a strong unidirectional multichannel waveform as posited by studies in the mouse (29). Second, we assessed whether the multichannel waveform of the NS-3 cluster was more likely to be symmetric in nature, especially compared to the other clusters, consistent with a stellate morphology (33).

The left panel of Fig. 6A shows the multichannel profile of an example neuron from NS-1 (with a narrow-spiking waveform) centered on the maximum amplitude channel which is generally assumed to be the channel closest to the soma. The right panel of the same figure shows the waveform as a function of time from each of the corresponding channels (see *Methods Section: Propagation Velocities*). The channels above the somatic waveform are more likely to have delayed peaks compared to the peak somatic waveform (shown in bold in Fig. 6A) suggesting a delay in the action potential as it travels upwards towards the pia.

**Figure 6:**
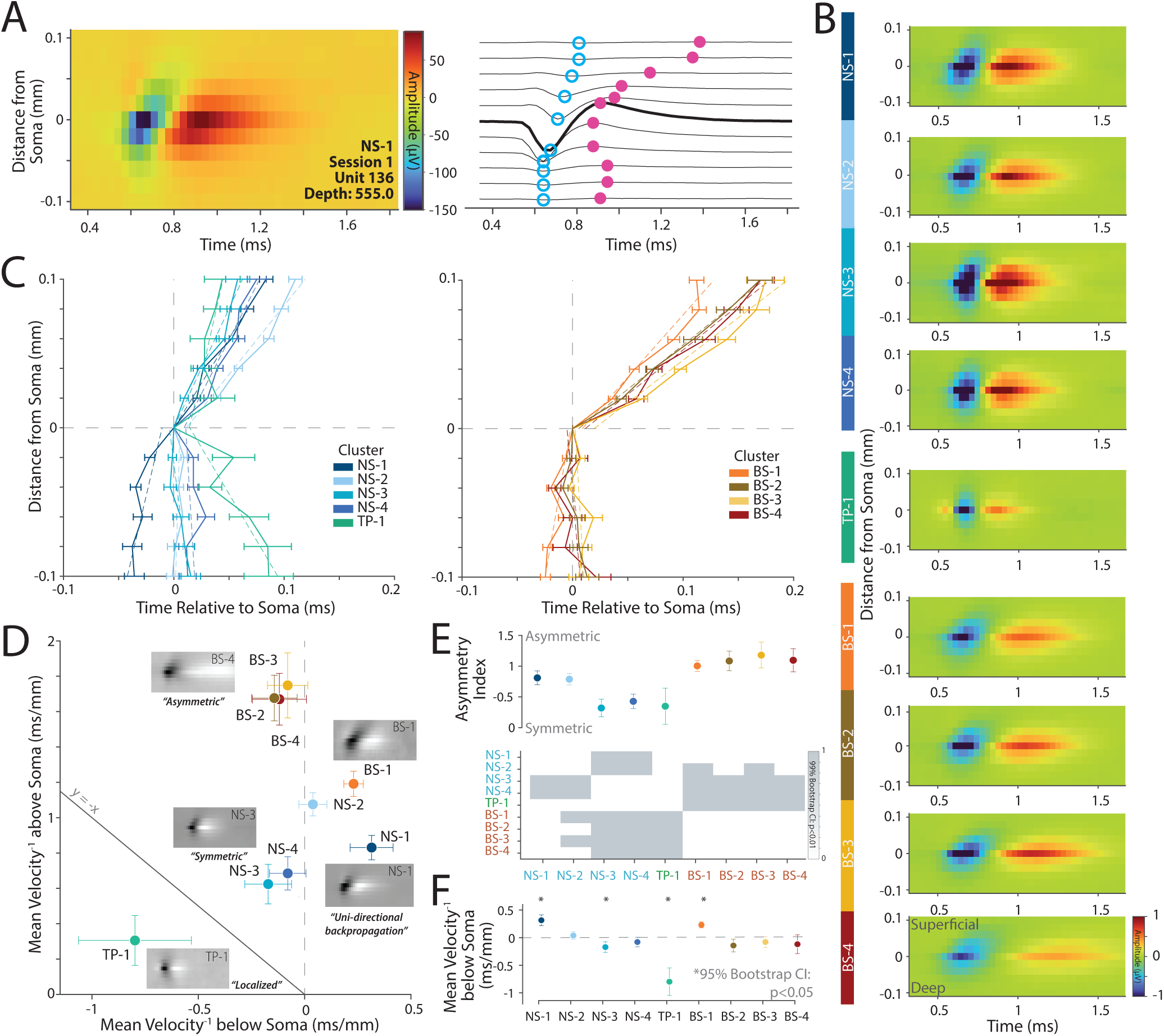
Multichannel waveforms reveal excitatory and inhibitory inclinations. (**A**)*Left*, A colormap of the maximum amplitude channel, assumed to be the position of the soma, and five channels above and below. The unit is identified by cluster, session, unit ID, and depth. *Right*, A time-series representation of each channel. The largest amplitude waveform is in bold. The time of waveform troughs (open blue points) and peaks (closed magenta points) of each channel are indicated. (**B**) Average multichannel extracellular waveforms per cluster. The colorbar shows the scale standardized for all clusters to map linearly between –1 and 1 *µ*v. (**C**) The average time of waveform trough at each channel location above and below soma (aligned to y-axis = 0, removing outliers using 95% confidence intervals) with standard error as errorbars. Velocities above and below soma and are estimated separately by linear regression (slope of the dashed line). *Left*, Narrow and Tri-phasic clusters. *Right*, Broad clusters. (**D**) Mean trough propagation velocity above soma vs below soma for each cluster. Clusters with positive slopes of propagation velocity both above and below the soma (unidirectional) are located on the right of x-axis = 0. Clusters which show positive slopes of propagation velocity above and negative below the soma (bidirectional) are located on the left of x-axis = 0, separated by degree of slope on the y-axis from the highest gradient in NS-3, to the lowest gradient in BS-4. Errorbars are standard deviation calculated from bootstrapped data (500 resamples). (**E**) *Top*, Asymmetry index of each cluster (see *Methods Section: Asymmetry Index*). This is the absolute value of the distance of each cluster’s propagation velocity above and below the soma, from the y=x line. *Bottom*, Errorbars are standard deviation calculated from bootstrapped data (500 resamples). An independent paired sample bootstrap was done to compare the asymmetry index of each cluster, shown in a heatmap with a value of 1 indicating a significant difference (99% bootstrap CI). (**F**) The mean slope of the propagation velocity below the soma was plotted per cluster. Errorbars are standard deviation calculated from bootstrapped data (500 resamples). An independent paired sample bootstrap was done to compare each slope to 0 as a measure of the propagation velocity direction with asterisks indicating significance (95% bootstrap CI).

Fig. 6B shows the average multichannel profile of the nine clusters. The BS clusters (BS1-4) show asymmetric waveforms with strong propagation of action potentials away from the the soma towards the pia. In contrast, NS-3 and NS-4 shows symmetric multichannel waveforms with minimal action-potential propagation away from the soma, whereas NS-1 propagates both to and away from the soma in the same direction. These visualizations are consistent with the predictions that NS-1 is more likely to be pyramidal in nature, whereas NS-3 is more likely to be stellate in nature.

We summarized the patterns in Fig. 6C by plotting the time of the trough relative to the distance from the soma (e.g., blue dots in Fig. 6A). The left plot shows the narrow-spiking and tri-phasic clusters and the right plot shows the broad-spiking clusters. On average, clusters NS-2 and BS2-4 show an asymmetric profile consistent with propagation above the soma towards the pia as predicted. In contrast, cluster NS-1 and BS-1 demonstrated *uni-directional propagation* consistent with morphologies of pyramidal neurons, whereas both NS-3 and NS-4 were far *more symmetric* in their multichannel profiles with only minimal propagation of action potentials away from the soma.

Fig. 6D shows a scatter plot of the propagation velocity above and below the soma for each cluster. We found that the different clusters fell into different quadrants on this plot. BS2-4 were largely asymmetric and had minimal propagation below the soma but strong propagation above the soma, whereas NS-2 was also modestly asymmetric. In contrast, NS-3 and NS-4 had more symmetric propagation above and below the soma. Finally, NS-1 and BS-1 showed unidirectional propagation both above and below the soma.

To quantify the symmetry or lack thereof for each cluster, we calculated an asymmetry index using the velocity of propagation above and below the soma (see *Methods Section: Asymmetry Index*, Fig. 6E), and also examined the velocity below the soma (Fig. 6F). Clusters NS-1, NS-2, and clusters BS1-4 had a large asymmetry index suggesting uneven propagation of the action potential away from the soma. In contrast, clusters NS-3, NS-4, and TP-1 were strongly symmetric, suggesting propagation of the action potential both above and below the soma. Symmetric action potentials are predicted for dense morphologies such as those observed for stellate neurons suggesting that NS-3 and NS-4 might possess stellate-like morphologies. Finally, NS-1 and BS-1 were the only clusters with a significant positive slope below the soma suggesting unidirectional propagation of the action potential (Fig. 6F).

Collectively, these results are consistent with our hypotheses that NS-1 is likely to be a excitatory neuron that is narrow-spiking in nature. In contrast, the multichannel waveform of NS-3 (and NS-4) is more consistent with dense dendritic morphologies around the soma, which is expected of non-pyramidal stellate neurons and thus consistent with anatomical descriptions of layer 4B neurons that project from V1 to MT (14).

### 2.6. Infragranular layers respond earlier to visual stimuli and precede activity in supragranular output layers

One of the key findings of V1 anatomy is the presence of strong connections from granular (layers 4A, 4B, 4C) and infragranular (layers 5/6) to supragranular (layers 2/3) layers (6). The functional prediction of this anatomical structure is that the responses of clusters in infragranular and granular layers should lead the clusters found in supragranular layers. A previous analyses of this dataset suggested this general pattern is true (26). Here, we further examined which candidate cell types contribute to this flow of information.

We first calculated the excitatory cross-correlograms (CCGs) of visually responsive neurons in all of these clusters (see *Methods Section: Cross-Correlations and Connectivity*). We normalized the CCGs by firing rate, jitter-corrected them to remove oscillations slower than 25 ms, and plotted significant pairs between clusters in a heatmap, sorted by cluster pairs. This analysis showed whether the reference cell type led or lagged the target cell type (Fig. 7A, B, Fig. S5A). Operationally, if the peak of the CCG occurs after 0, this indicates the reference cell is leading the target cell, whereas a peak in the CCG occurring before 0 indicates the reference cell class is lagging the target cell class. To measure this directionality, we calculated a lead-lag index for each significant CCG by subtracting the CCG values on the right side of time lag 0 with CCG values on the left side of time lag 0, and then normalized by their sum. Therefore, if a reference cell is leading a target cell, the index will be 1.

**Figure 7:**
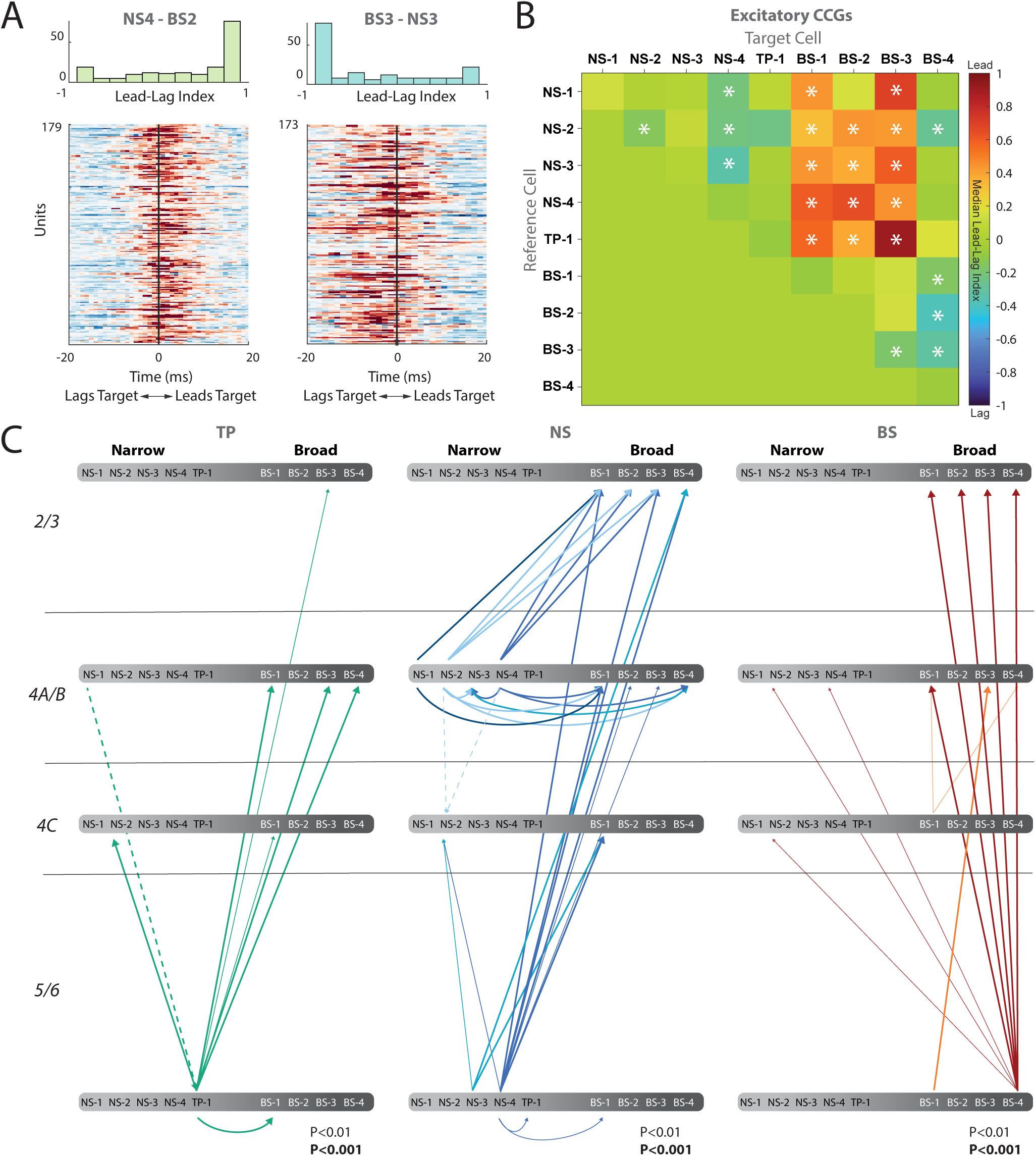
Cross-correlations show mostly feedforward lead-lag connectivity between clusters and among layers of V1. (**A**) Example Cross-correlograms (CCG) between clusters. *Top*, Histograms of CCG asymmetry. *Bottom*, Heatmap of pairwise units latencies. *Left*, Cluster NS-4 leading BS-2. *Right*, Cluster BS-3 lagging NS-3. (**B**) Heatmap of pairwise cross-correlation median Lead-Lag indices between clusters which excite each other. The reference cell (left) cluster interacts with the target cell (bottom). If the reference leads the target, the color is red, if it lags behind the target, the color is blue. * indicates Wilcoxon signed rank sum test significance (p < 0.05). (**C**) CCGs between clusters and layers, separated by the tri-phasic (TP), narrow (NS), and broad-spiking (BS) groups. Arrows indicate directionality of CCG, from lead to lag of significant interactions between cluster and layer. Bold arrows indicate p < 0.001, Small arrows indicate p < 0.01. Solid lines indicate feedforward directionality, dashed lines indicate feedback directionality.

Fig. 7A shows example CCGs and the lead-lag index for NS4-BS2 (left panel) and BS3-NS3 (middle panel). The reference cell class NS-4 leads the target cell class BS-2, whereas BS-3 lags the NS-3 class. These examples suggest that the NS clusters are more likely to lead the BS clusters. We assessed significance of this index by using a sign-rank test and summarized all the pairwise relationships using a colormap (Fig. 7B, see *Methods Section: Lead-Lag Index*). These analyses revealed that narrow-spiking clusters largely lead the broad-spiking clusters BS1-3, with the exception of BS-4. This BS-4 cluster also led all other broadspiking clusters, as well as cluster NS-2. Moreover, within the narrow-spiking clusters, NS-4 noticeably led NS-1, NS-2, and NS-3 clusters.

The colormap in Fig. 7B reveals broad relationships between clusters but does not incorporate layer information. We further quantified and summarize these lead-lag relationships between different pairs of candidate cell types and layer. Fig. 7C plots a directed graph with the line thickness indicating the strength of the relationship (estimated from the p-values: thin lines p < 0.01, and thick lines p < 0.001, Fig. S5A). Our analysis of all significant units between pairs of clusters shows a general trend of narrow-spiking clusters (NS-1, NS-2, NS-3, NS-4, and TP-1) in layer 4 leading broad-spiking clusters (BS-1, BS-2, and BS-3) in layers 2/3 with the exception of BS-4, largely localized to layers 5/6, which also leads the other broad clusters (Fig. 7B). Together, significant interactions between clusters and among layers (Fig. 7C) suggest that there is a flow of information from infragranular and granular layers to supragranular layers. This is consistent with the feed forward model of local circuitry suggested anatomically (6).

Finally, we computed the inhibitory CCGs between pairs of neurons within a cluster. While there were far fewer significant pairs, NS-3 and NS-4 show a center peak (Fig. S5B), with inhibitory troughs on each side, suggesting the underlying circuitry may have a strong common excitatory input as proposed in analytical studies (51).

### 2.7. Tri-phasic waveforms and positive spikes likely emerge from return currents and axons

Our study thus far has largely focused on the interesting links between functional and structural properties of the biphasic negative waveforms (NS and BS groups). Our WaveMAP analysis also revealed extremely narrow tri-phasic waveforms (TP-1, with trough-peak duration = ~170 *µ*s) with very average functional responses, across the board. These waveforms were found both in gray matter in layers 4A, B, C through 6 as well as in white matter (Fig. S4B, D). For this TP-1 class, direction or orientation selectivity was largely uniformly distributed and unremarkable.

To better understand this TP-1 neural population, we examined the multichannel waveform of the TP-1 type and found two main types of profiles. The first, TP type A, had a localized multichannel waveform profile (Fig. S6A) and was present throughout cortex and white matter and was largely uniformly selective for orientation and direction. These neurons with a small and local multichannel waveform are likely balancing return currents when a spike is generated in the axon initial segment (42). The second type, TP type B, shows a much larger spread across channels (Fig. S6A), and these were mainly located within layers 4A/B, 4C and 5/6 of cortex. They also had lower direction and orientation selectivity (Fig. S6B). The larger spread exhibited by TP Type B is reminiscent of recordings with Neuropixels along a retinal ganglion cell axon in the superior colliculus of the mouse (52). Thus, the multichannel subtype B of the TP-1 group is consistent with propagation through a neurite aligned vertically, or close to the axon initial segment.

We also found that positive spiking waveforms were quite common in the Neuropixels recordings. The majority of these positive spiking units were found in the white matter, with a smaller fraction more likely in layers 5/6 (84%, *χ*^2^ (1, 95) = 41.779, p *<* 0.0001, Fig. 1F). WaveMAP clustering on these waveforms revealed five clusters (Fig. S7A), all concentrated in white matter and deep layers 5/6 (Fig. S7B). We investigated the multichannel profiles of these neurons and observed a large spread across channels similar to TP type B (Fig. S7C). Functionally, these positive spiking neurons were largely unresponsive to visual stimulus (only 32/105 responsive units) and did not demonstrate any remarkable responses to the visual stimuli (Fig. S7D, E). Collectively, these results suggest that these positive spiking waveforms are likely axons passing through the white matter (42).

## 3. Discussion

The goal of this study was to analyze the relationship between structure and function and thus better understand the microcircuit in monkey V1. To this end, we reanalyzed high-resolution Neuropixels recordings across layers of V1 of two anesthetized rhesus macaques while we presented drifting Gabor gratings in 36 orientations (26). We used our non-linear dimensionality reduction and clustering approach (WaveMAP, 27, 28, 38, 39) to group the diversity of waveform shapes collected in these large neuron population recordings from these negative spiking units into 9 putative cell classes. Candidate cell classes also exhibited distinct functional properties with respect to their laminar organization, tuning properties, bursting patterns, and multichannel waveforms. Within these candidate cell classes, we discovered links between structure and function that have not been reported.

One of the key findings from this study is better identification of the layers in which we found different candidate cell types. While some studies have shown improved cortical layer identification using higher frequency band signals (53), we used CSD in combination with histological estimates to determine the layer boundaries of our recordings. Such analyses revealed that narrow-spiking neurons were approximately 30% of the neurons that we recorded in V1, and > 75% of these neurons were found in layers 4A/B, and 4C. These *in vivo* functional findings are remarkably consistent with reports from anatomical studies that 80% of Kv3 and especially Kv3.1b positive neurons are found in layers 4A/B and 4C of V1 (13).

However, this ~ 30% proportion is much larger than what is expected from anatomical studies. It was reported that only ~ 7% of all neurons in a V1 column are Kv3.1b positive and thus likely to be narrow-spiking (13). This discrepancy between our findings and the anatomical studies could emerge for several reasons. First, WaveMAP is an unsupervised clustering method: it is possible that we have overestimated the fraction of narrow-spiking neurons. Even the narrow-spiking neurons that we identified had a range of spikewidths (0.27 ms and 0.30 ms, for NS-1 and NS-2 classes, and 0.2 to 0.24 ms for NS-3 and NS-4 classes). One possibility is that the Kv3.1b positive units are only the NS-3 and NS-4 classes which had widths closer to 0.2 ms, in which case the numbers would be more similar to what is observed in anatomical and in vitro studies (13, 54, 55). Second, that particular study (13) only identified the number of Kv3.1b positive neurons of V1 compared to the overall number of cells. However, there are also other potassium channel types of the Kv3 family (such as Kv3.2), which can also facilitate narrow-spiking and these are known to be expressed in V1 (12), and other brain areas of primates (56). Further assessment of the expression levels of these voltage-gated potassium channels and sodium channels that can confer fast spiking will help better estimate the population of fast-spiking cells in V1, which in turn will inform physiological studies (57). Third, we don’t know what fraction of any given cell type is active during *in vivo* recordings (58). Recordings in auditory cortex of anesthetized rats suggest that layers 4A/B, and 4C neurons (an input and feedforward layer) are more active during anesthesia compared to neurons in layers 2/3 and layers 5/6 (feedback layers) (59), which would help explain the over representation of narrow-spiking neurons in our dataset.

We identified four narrow-spiking clusters with distinct properties. Do any of the clusters map onto a parvalbumin (PV) specific inhibitory population? Anatomical studies of V1 suggest that PV neurons are observed in all layers, but are more likely in layers 4A, 4B, and 4C. Currently, there are no functional studies from monkey V1 of PV neurons with ground truth optotagging. However, *in vivo* mouse studies suggest that PV neurons in V1 have narrow, small amplitude waveforms with symmetric forward and back-propagation profiles (33). Based on these observations, we suggest NS-4 (and likely the small amplitude neurons in cluster NS-3 not located in layers 4A/B) are likely to be PV neurons. This assertion is based on five lines of evidence. First, NS-4 neurons were found throughout all the layers of cortex. Second, they also were one of the narrowest clusters observed with a mean trough to peak width of 0.24 ms. Third, they had symmetric multichannel waveforms as expected of aspiny cells with dense dendritic and axonal morphologies around the soma (29, 60). Fourth, they responded early after stimulus onset and had robust firing rates to stimulus onset that was earlier than all the broad-spiking neurons (61). Fifth, cross correlation analyses suggested NS-4 neurons often have cross correlograms that suggest that these neurons mutually inhibit each other, consistent with the autaptic and mutually inhibitory connections between these neurons (62), which can lead to mutual suppression of inhibitory neurons (Fig. S5B). Again, rigorous optotagging techniques (63) with inhibitory neuron specific viral constructs (e.g., those with mDlx 2.0 or other enhancers (64)) combined with Neuropixels recordings in monkey V1 will help establish whether PV neurons show the same layer distribution, and waveform properties as NS-4.

We identified *in vivo* a population of neurons that are highly-direction selective, localized to layer 4B, and possessing multichannel extracellular waveforms that are consistent with those associated with stellate morphologies. We speculate that these neurons are likely neurons that project from V1 to MT. These neurons had narrow-waveforms with a trough to peak width of 0.2 ms. The prediction for anatomical studies is that these MT projecting V1 neurons will likely express a fast-spiking potassium or sodium channel. Intriguingly, one of the groups of Kv3-immunoreactive neuronal populations in anatomical studies of V1 had large, elongated cell bodies and the study suggests that these are likely “spiny stellate cells with horizontally extended dendritic fields in layer 4B” (12). However, due to incomplete staining of cell arbors, the authors could not conclusively classify these cells as stellate. We suggest that experiments that use tracers (e.g., non-toxic rabies virus) to identify neurons in V1 that project to MT (14) with approaches for examining ion channel expression could help test this prediction more rigorously (13).

We found that ISI distributions of the recorded cell classes were highly diverse. Some neurons demonstrated classical broad ISI distributions with a slow decay consistent with a Poisson process and a refractory period. However, many other neurons showed ISI distributions that had a strong peak within 10 ms. The typical assumption is that cortical neurons are largely Poisson in nature (65). However, our results suggest that even at the earliest stages of visual processing, many neurons can show ISI distributions that are not fully consistent with the Poisson assumption. We have interpreted these neurons with peaky ISI distributions as “bursting”. Our results are consistent with the findings that most of the neurons in area V1 and MT burst in response to a sensory stimulus (50, 66). Our results also reaffirm reports of bursting in anesthetized monkey V1 (49), and more recent studies suggesting the presence of narrow spiking, bursting, excitatory neurons in awake monkey V1 (19). A key advance of this study over these previous studies is that we also identified the laminar location of these neurons and showed that superficial layers and in layers 4A/B and 4C are more likely to burst than neurons in deeper layers. One caveat to these findings of bursting in some populations is that we used isoflurane anesthesia which could potentially alter the network properties such as neurotransmitter release and impact inter-spike intervals in these neurons (67–69) and thus our findings need to be further validated in awake animals.

The unique advantage of high-resolution laminar recordings with Neuropixels over other approaches for recording spiking activity *in vivo* is that it densely samples the electrical field around neurons leading to a detailed spatiotemporal profile of the extracellular waveform for each neuron. Guided by previous efforts (29, 33), we analyzed how the waveforms vary across electrodes and derived various metrics including the spread and propagation velocity for each of these clusters. These metrics could provide additional information to separate putative cell classes and understand how the morphology of a neuron might affect high-density extracellular recordings. For instance, at least for the mouse, clustering on multiple modalities such as inter-spike-interval, waveform shape, autocorrelogram, and features enables better separation of cell classes that are closer to ground-truth (70).

The multichannel waveform contains useful information about the morphology of the units we record from, as well as properties of the cortex. Considering that in V1, most neurons show a propagation above the soma towards the pia. This feature can help us understand the layout of deep cortices. While it may be trivial for superficial V1 recordings, with recordings made deep in the cortex, it is often hard to tell where the surface of the second cortex is, or how perpendicular the probe is. In our Sessions 4-5, the waveform trajectories were inverse, and had to be flipped due to the layout of the cortex underneath. Additionally, the velocity of the propagation could possibly tell how perpendicular a recording is (33). For example, if the velocity and spread are bigger for perpendicular recordings compared to more tangential recordings. While we did not observe these differences in this dataset, it may be an important point to consider for future analyses.

Analyses of the multichannel waveform also helped to understand the diversity within the narrow-spiking group of cell clusters that was not obvious in the single channel waveform Fig. S6C. We showed that NS-1, a candidate cell class with ISI distributions consistent with previous reports of bursting, had strong unidirectional propagation towards the soma suggesting a particular morphology for these neurons. In contrast, we found that NS-3 and NS-4 clusters had strong symmetric spatiotemporal profiles around the soma suggesting that their morphology was different from the NS-1 cluster and more consistent with dense local dendritic arborization. The NS-3 and NS-4 clusters also had the largest spiking amplitudes. Modeling studies suggest that dense local dendritic arborization exerts strong effects on the amplitude of the extracellular waveform but minimal to no effects on the shape of the waveform (33). We also found that the BS-1 cluster had multichannel properties similar to those observed for NS-1, with a near unidirectional propagation away from the soma. However, the BS-1 cluster also had the smallest spike amplitude, and did not share the laminar compartments nor functional properties of NS-1. In contrast, the multichannel waveforms of BS clusters 2-4 largely propagated away from the soma toward the pia and had minimal to no propagation towards the white matter. BS2-4 were heavily localized in layers 4C and 5/6. Anatomical studies of layer 4C stellate cells suggest a strong axonal arborization towards layers 2/3 (71). Similarly, both dendritic fields and axon paths of many layers 5/6 neurons are found in supragranular and infragranular layers (72). We believe such morphologies are likely to show more propagation of activity away from the soma.

We used WaveMAP, our novel non-linear dimensionality reduction approach, to cluster waveforms into candidate clusters and identify underlying diversity in high-density Neuropixels recordings. Our approach improves on classical efforts for identifying cell types from extracellular recordings that only rely on the width of the waveform or a few features to separate cell populations into narrow and broad-spiking neurons. Nevertheless, the clusters identified are candidate cell classes and need validation with rigorous ground truth optotagging (73). Fortunately, work from the Allen Institute and others has finally led to AAV viral constructs with promoters specific for inhibitory neurons (64, 74, 75), and projection targets *in vivo* (76). We have recently used these constructs in anesthetized animals and reliably optotagged inhibitory neurons in premotor and prefrontal cortex (63). Combining optotagging with the approaches here might enable even better delineation of candidate cell types *in vivo*. Finally, recently available high-density Neuropixels Ultra may enable more precise localization of neurons, cleaner signal-to-noise ratios, and better estimates of multichannel waveforms in these brain areas (77). Combining such approaches and careful analyses as performed here will enable a more detailed understanding of the microcircuit in V1.

## 4. Methods

Several methods sections are adapted from Trepka et al. (2022)(26) as the same dataset is reanalyzed in this study but with a focus on candidate cell types and their properties. For completeness and readability, we briefly replicate some of these methodological details here. The majority of the methods focuses on key details about the WaveMAP approach and functional assessments.

### 4.1. Subjects

The dataset used in this paper is composed of electrophysiological recordings in 2 anesthetized adult male rhesus macaques (Macaca Mulatta, M1, 13 kg; M2, 8 kg). Experimental data were collected under procedures that were designed in full accordance with National Institutes of Health Guide for the Care and Use of Laboratory Animals, the Society for Neuroscience Guidelines and Policies. All experimental and surgical procedures were approved by the Institutional Animal Care and Use Committee (IACUC) protocol (APLAC-9900) of Stanford University.

### 4.2. Surgical Details and Electrophysiological Recordings

#### 4.2.1. Surgical Details

Our workflow for the surgery and electrophysiological recordings were as follows. 24 hours before the recording session, the monkey was treated with dexamethasone phosphate to reduce cerebral edema. Then, on the day of the recordings, we sedated the animal using ketamine HCl (10 mg/kg body weight, intramuscularly). Monkeys were then intubated and ventilated with 1-2% isoflurane in a 1:1 mixture of nitrous oxide and oxygen to maintain general anesthesia, and placed in a stereotaxic frame. During the surgery, we monitored electrocardiogram, respiratory rate, body temperature, blood oxygenation, end-tidal CO_2_, urine output and inspired/expired concentrations of anesthetic gases to ensure stable anesthesia. Normal saline was administered intravenously at a variable rate to maintain adequate urine output. Once animals were stable on the anesthetic plane, we administered a cycloplegic agent (1% atropine sulfate for pupil dilation) and focused the eyes with contact lenses on an LCD monitor. Finally, we used vecuronium bromide (60 *µ*g/kg/hr) to prevent eye movements.

#### 4.2.2. Electrophysiological Recordings

We first performed an occipital craniotomy over the opercular surface of V1 and then reflected the dura to expose a small (~3 *mm*^2^) patch of cortex. Next, we identified a region relatively devoid of large surface vessels and inserted the Neuropixels probe with the aid of a surgical microscope. The Neuropixels 1.0 probe is quite thin (70 *µ*m x 20 *µ*m) and sometimes flexed upon contacting the pia. Thus, insertion sometimes required multiple attempts if it flexed upon contacting the pia. The junction of the probe tip and the pia could be visualized via the (Zeiss) surgical scope and the relaxation of pia dimpling was used to indicate penetration, after which the probe was lowered at least 3–4 mm. Prior to probe insertion, we dipped the Neuropixels probe in a solution of the DiI derivative (FM1-43FX, Molecular Probes, Inc.). The dye was used for subsequent histological visualization of the electrode track (e.g., Fig. S1).

The Neuropixels 1.0 probe has 986 contacts throughout the length of the probe (1 cm), of which 384 can be selected at any given point for recordings. We selected either the opercular surface cortex (M1) or within the underlying calcarine sulcus (M2). Recordings were made at 1–3 sites in one hemisphere of each monkey. At the end of the experiment, monkeys were euthanized with an overdose of pentobarbital (150 mg/kg) and perfused with normal saline followed by 1 liter of 1% (wt/vol) paraformaldehyde in 0.1 M phosphate buffer, pH 7.4.

#### 4.2.3. Visual stimulation

Visual stimuli were presented on a LCD monitor NEC-4010 (88.5 (H)* 49.7 (V) cm, 1360 × 768 pixels and a frame rate of 60 Hz) positioned 114 cm from the monkey. We first identified receptive fields (RFs) from online multi-unit activity and localized on the display using at least one eye. RF eccentricities were ∼4–6^◦^ (M1) and ∼6–10^◦^ (M2). We then presented circular drifting Gabor gratings (2^◦^/sec., 100% Michelson contrast, 1.5 degrees of visual angle in diameter) within the joint RFs of recorded neurons either monocularly or binocularly. Gratings drifted in 36 different directions between 0–360° in 10^◦^ steps in a pseudorandom order. We used four spatial frequencies (SF, 0.5, 1, 2, 4 cycle/^◦^). Optimal SFs and eye conditions were determined offline to categorize V1 neurons into simple or complex neurons. We used a stimulus duration of 1 second and repeated it 5 or 10 times and presented a blank screen with an equal luminance to the Gabor patch during the inter-stimulus interval (0.25 s).

### 4.3. Scaling Laminar Depths

We estimated the depth of each unit relative to the boundary between layers 4C and 5/6, using the current source density analysis and histology results identifying the electrode tracks. This process is outlined in more detail in Trepka et al. (2022) (26), and we provide a brief description here. For each recording, we first performed the current source density (CSD) analysis on the stimulus-triggered average of LFP. LFP signals recorded from each of the 4 neighboring channels were averaged and realigned to the onset of the visual stimulus. CSD was estimated as the second-order derivatives of signals along the probe axis using the common five-point formula (78). The result was then smoothed across space (*σ* = 120 *µ*m) to reduce the artifact caused by differences in electrode impedance. We located the lower boundary of the major sink (the reversal point of sink and source) as the border between layers 4C and 5/6. Based on this anchor point, we assign other laminar compartment borders using histological estimates. Although in V1 layers 2, 3, 4A, 4B, 4C*α*, 4C*β*, 5, and 6 are distinct layers, we combine layers 2 and 3, 4A and 4B, 4C*α* and 4C*β*, and 5 and 6 for analysis.

For the scaling of each session, the layer boundaries of all five sessions were averaged to obtain a common reference of laminar organization. For each session, the units within each layer assignment were scaled to their equivalent depths within the average boundary assignment. In sessions 4 and 5 (M2), the probe recorded from the calcarine sulcus, and all depths were inverted before scaling. It is important to note the experimental differences between sessions, including that sessions 4 and 5 did not have many units recorded from layers 2/3 (n = 5 units), and session 2 showed minor drift in its receptive field properties (25). However, our functional analyses of the receptive field properties suggests that all 5 recording sessions were within the same area and largely perpendicular to the surface of cortex (Fig. S1H, Fig. S3A, B, D).

### 4.4. Spike Sorting and Data Curation

We used Kilosort 2 to extract templates of the extracellular waveform of each unit (35). To convert Kilosort 2 template projection amplitudes to a good estimation of raw amplitude, a scaling factor from the acquisition system of 2.3 was applied for units of *µ*V. After spike sorting, we isolated the maximum amplitude channel from the template for each unit, as well as 5 channels above and 5 below the maximum amplitude channel for the multichannel trajectory (see below). Second, each maximum amplitude waveform was then normalized between 1 and –1. Finally, we separated the dataset into positive and negative spiking units, with the maximum peak of positive units occurring before the minimum trough.

In our past experience, we have found that rigorous and conservative spike sorting is a prerequisite for delineating cell types using WaveMAP (28). To obtain high-quality single units that could then be passed to WaveMAP, we used the following semi-automatic quality control method. This method involved two parallel steps. In the parallel step 1, we developed a graphical user interface (GUI) to manually curate all the units. The GUI presented the normalized maximum amplitude waveform, as well as three user selectable annotations: “good”, “noisy”, and “artifact”. For this step, two reviewers (NC and AP) curated all 2,529 templates from all 5 sessions in the first pass. The waveforms that were rated “good” by both reviewers were confirmed, and then any units that received a single “good” label or “noisy” by either reviewer were run through a second pass of inspection. Finally, we only chose units that were annotated as “good” by both reviewers. We calculated both internal consistency through Chronbach’s Alpha (*α* = 0.72), and the inter-rater reliability through Cohen’s Kappa (*κ* = 0.69), which gave us acceptable agreement on internal consistency, as well as substantial agreement for inter-rater reliability. In the parallel step 2, we filtered units by SNR and excluded units with SNR thresholds below 0 or above 3.7 (95% CI Range: [0.3350 1.0641]). The final dataset was the intersection of the units that passed both the manual curation step and the SNR threshold. After this quality control, our final dataset (regardless of visual responses) with positive and negative spiking waveforms was composed of 905 units.

### 4.5. WaveMAP Analysis

We applied our novel WaveMAP approach (27, 28) to our dataset, with positive and negative units separated to prioritize local over global clustering. We first applied UMAP to the normalized maximum amplitude waveforms after spike sorting and quality control to obtain a graph, and then passed this graph to Louvain clustering to delineate clusters (N_neighbors = 20; MIN_DIST = 0.2; RESOLUTION = 1.0). More technical details of the WaveMAP approach are available in the original publication (27). The 801 negative spiking units were partitioned into 9 clusters. The 104 positive spiking units were partitioned into 5 clusters (Fig. S7A).

We also used a classification approach to assess whether the identified clusters were well separated from each other, and found that it was indeed the case (88% standard accuracy over 9 clusters), which suggests we did not overcluster on this dataset (Fig. 2C). In addition, the Louvain clustering approach is a hierarchical approach, and so higher resolution parameters lead to increased importance on global vs. local features, and larger and fewer clusters. Thus, when applying a higher UMAP resolution parameter (RESOLUTION = 2.5), our narrow-spiking neurons combined into one large cluster showing that the putative cell classes in the narrow-spiking group are closer to each other in high-dimensional space than to the other groups (Fig. 2D). We also used an alternative clustering approach termed ensemble clustering for graphs (ECG, 37)), a consensus clustering method that simultaneously evaluates various resolution parameters, and we found that this clustering also resulted in nine candidate cell types and largely overlapped with our UMAP approach (Fig. 2E).

To better understand why WaveMAP separated these waveforms into these clusters, we performed three classical operations on the waveforms of this dataset. First, we calculated classical features of the waveform, including the trough-to-peak width, or the time between trough and peak, the trough-to-peak amplitude, which is the raw voltage change between the trough and the peak, and the repolarization time, defined as the time between the peak and 1/2 the peak of the repolarization curve (Fig. S2A). We then scaled the size of each unit point in UMAP space by the width of the waveform, the time for repolarization, and the amplitude (Fig. S2B-G). This plot revealed that both UMAP dimension 1 and 2 were strongly correlated with both waveform duration and repolarization time. An important step in pre-processing waveforms for WaveMAP is to normalize the waveforms so amplitude should not factor in the clustering. Nevertheless, when we scale the size of the markers of each unit by its relative amplitude, we found that WaveMAP identifies that some clusters have larger amplitudes than others, again highlighting that our approach can separate clusters with biologically relevant differences.

### 4.6. Single Neuron Response Properties

To characterize neuronal properties, the evoked activity was assessed using mean firing rate (spikes/sec) over the whole stimulus presentation period, offset by response latency delay. Only responses to the preferred spatial frequency and eye conditions were selected. The maximum firing rate was the neuron’s response to the preferred drifting orientation and direction.

#### 4.6.1. Receptive Field Properties

We assessed the following receptive field properties. These properties have been used extensively in past literature to characterize V1 neurons *in vivo* and provide a robust assessment of functional properties of V1 neurons.

1. Direction selectivity (Direction Index, DI) was determined as the response to preferred orientation and drift direction minus the response to preferred orientation but opposite drift direction, divided by the sum of these two responses (79). 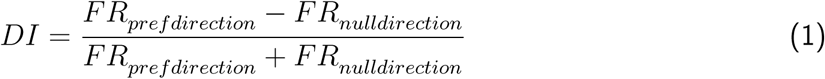
2. Orientation selectivity (Orientation Index, OI) was determined as the response to preferred orientation minus the response to orthogonal orientation, divided by the sum of these two responses (79). 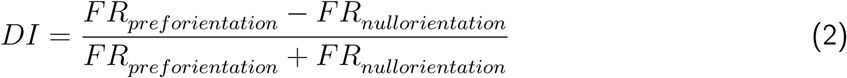
3. Simple Complex Index, also referred to as Modulation Ratio in prior studies, was defined as F1/F0, where F1 and F0 are the amplitude of the first harmonic at the temporal frequency of drifting grating and constant component of the Fourier spectrum to the neuron’s response to preferred orientation. A simple-complex index of less than 1 was assigned as complex cell, and greater than 1 as a simple cell (20, 80).

#### 4.6.2. Peri-stimulus time histogram – PSTH

For each unit, we estimated a trial-averaged peri-stimulus time histogram (PSTH) for the preferred stimulus orientation and direction as the firing rate in spikes/s between the period of 50 ms before and 500 ms after stimulation. 67 ms were added to the stimulation time to correct for the time for the screen to update the visual stimulus. We then Gaussian smoothed with a 13 ms kernel. The preferred stimulus was determined by selecting the condition with the maximum firing rate in the stimulus period. The standard error of the mean PSTH was calculated for each unit. For each cluster, an average PSTH included the trial-averaged PSTH of each unit in that cluster. For each layer, an average of the PSTHs of all units within the bounds of the layer was calculated.

#### 4.6.3. Inter-spike Interval

We estimated the burstiness of the units by analyzing the inter-spike interval (ISI) distribution. The inter-spike interval was estimated using the difference in all spike times per unit throughout the entire recording session, with 0.4 ms bins. We chose this bin size as it most clearly represented the structure of the ISI density distributions, without overemphasizing random noise or artifacts. The ISI histograms were normalized to a relative probability, where the number of elements in each bin relative to the total number of elements in that bin is 1 at maximum. Visual inspection suggested considerable heterogeneity in these ISI distributions.

#### 4.6.4. Bursting Index

To analyze this heterogeneity, we used principal component analysis (PCA) to determine the various components of these ISI distributions. We then estimated the principal component coefficients for each unit. We plot the mean loadings of each cluster and their standard error of the top three principal components against each other. The difference between the first and third PC loadings make up our bursting index, which we plot per unit against laminar depth and orientation selectivity. Finally, we plot the median orientation index and bursting index per cluster against each other, and performed a shuffle test (500 shuffles) for both the bursting and orientation indices to determine the 2.5 and 97.5 percentiles of the shuffled distribution.

### 4.7. Multichannel Profiles

The extracellular action potential waveforms from five channels above and five channels below the maximum amplitude channel were extracted to build the multichannel profile per unit. Only the linear column of channels to which the maximum amplitude unit waveform belonged was plotted for ease of visualization, leaving eleven channels for analysis. A color map of the channel activity shows the 3D representation of the neuronal activity during an action potential, averaged for each cluster. The color bar is scaled the same for each cluster, to give a clear indication of both the shape and overall amplitude.

#### 4.7.1. Propagation Velocities

A time-domain analysis of the troughs of the multichannel profile shows mean propagation trajectories of each cluster (only including the 95th percentile of all trough times to exclude outliers), with errorbars showing standard error of the mean trough time per cluster at each channel. We used linear regression to calculate the slope of the propagation above and below the soma, which are referred to as propagation velocities (29). The propagation velocity is reported in units of (ms/mm), and so is also referred to the inverse velocity (velocity^−1^). We plotted the mean propagation velocity above and below the soma against each other (velocity^−1^*_below_*, velocity^−1^*_above_*) to gain a metric of velocity symmetry around the soma. We used bootstrapped trough times (500 resamples) for each cluster to find the standard deviation of the propagation velocities per cluster. On the velocity^−1^*_below_* axis, if a cluster is located higher than 0, the slope below the soma continues in the same direction as above the some, meaning the propagation velocity is unidirectional. If it is less than 0, the velocity slope below the soma is in the opposing direction than the slope above the soma, meaning the propagation velocity is bidirectional. We tested which clusters showed a significant difference in propagation velocity below the soma from 0 using an independent paired sample bootstrap test (95% Bootstrap CI: p *>* 0.05). On the velocity^−1^*_above_* axis, the slope increases roughly in the order of peak-trough duration.

#### 4.7.2. Asymmetry Index

We calculated the index of symmetry (SI) as the orthogonal distance between each point (velocity^−1^*_below_*, velocity^−1^*_above_*) and the diagonal line y = –x (33). The equation is as follows:

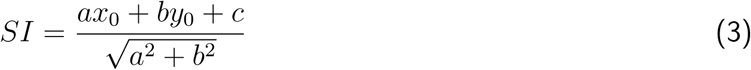

where (*x*_0_,*y*_0_) = (velocity^−1^*_below_*, velocity^−1^*_above_*), a = 1, b = 1, and c = 0 for y = –x. An unsigned asymmetry index was then calculated, which does not include information on which direction (above vs. below the soma) was dominant. A small value of the asymmetry index would indicate more symmetry between the velocity slopes above and below the soma, whereas a larger value indicates more asymmetry. A negative value indicates that the velocity slope below the soma is greater than the slope above the soma. We calculated a paired sample independent bootstrap test for significant differences between all clusters, plotted in a heatmap where a value of 1 indicates a significant difference between pairs (99% Bootstrap CI: p *>* 0.01).

### 4.8. Cross-Correlations and Connectivity

Cross-correlations between spike trains of all pairs of simultaneously recorded neurons were computed (26). Spiking activity was chosen within the 0.4–1 s window of each visual stimulus presentation. The raw cross-correlogram (CCG) for a pair of neurons (j,k)was defined as follows:

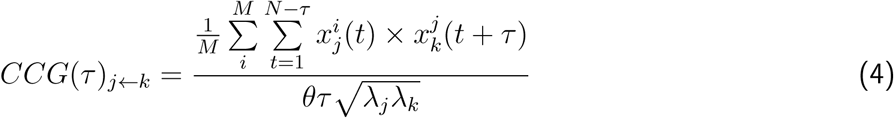

where *M* is the number of trials, *N* is the number of time bins within a trial, *τ* is the time lag, *x_ij_*(*t*) is one if neuron j fired in time bin t of trial i and zero otherwise, and *λ_j_* is the mean firing rate of neuron j computed over the same bins used to compute the CCG at each time lag. *θ*(*τ*) is a triangular function, *θ*(*τ*) = *N τ*, that corrects for the difference in the number of overlapping bins at different time lags. We denote the CCG computed with neuron j as the first (reference) neuron and k as the second (target) neuron in the correlation function as j-k.

Raw cross-correlograms (CCGs) were normalized by the geometric mean of firing rates of the pairs and then jitter-corrected to remove the fluctuations slower than 25ms. For excitatory interactions, a CCG was determined significant if its peak occurred within 10 ms of zero-time lag, and its peak value exceeded 7 standard deviations above the mean of the noise distribution (defined as CCG values of 50-100 ms from zero-time lag). For inhibitory interactions, a CCG was considered significant if its trough occurred within 10ms of zero-time lag, and its trough value fell below 5 standard deviations from the mean of the noise distribution.

#### 4.8.1. Lead-Lag Index

The lead-lag index is a measure of correlogram asymmetry (CA), calculated as the values on the right (lead) side of the CCG, subtracted by the left (lag) side, and then normalized by their sum (e.g., a value closer to 1 means that the reference cell is leading the target cell). If the CA is within [-0.3, 0.3], it is assumed that the CCG is more symmetrical, reflecting a common input, whereas a CA larger than 0.4 or smaller than –0.4 may reflect synaptic connections (81).

## 5. Acknowledgments

We thank Jonathan C Horton for extensive help with the recordings and histology. We thank Tim Harris and Karel Svoboda for providing the Neuropixels probes, Shellie Hyde and Sam Baker for technical assistance.

## 5.1. Funding

We thank the following institutions for providing funding necessary for the completion of this work:

Whitehall Foundation 2019-12-77 (CC)

National Institute of Neurological Disorders and Stroke K99/R00NS092972 (CC)

National Institute of Neurological Disorders and Stroke R01NS122969 (CC)

National Institute of Neurological Disorders and Stroke R21NS135361 (CC)

National Institute of Neurological Disorders and Stroke NS116623 (TM)

National Eye Institute EY014924 (TM)

National Eye Institute EY029759 (TM)

Howard Hughes Medical Institute (TM)

## 5.2. Author Contributions

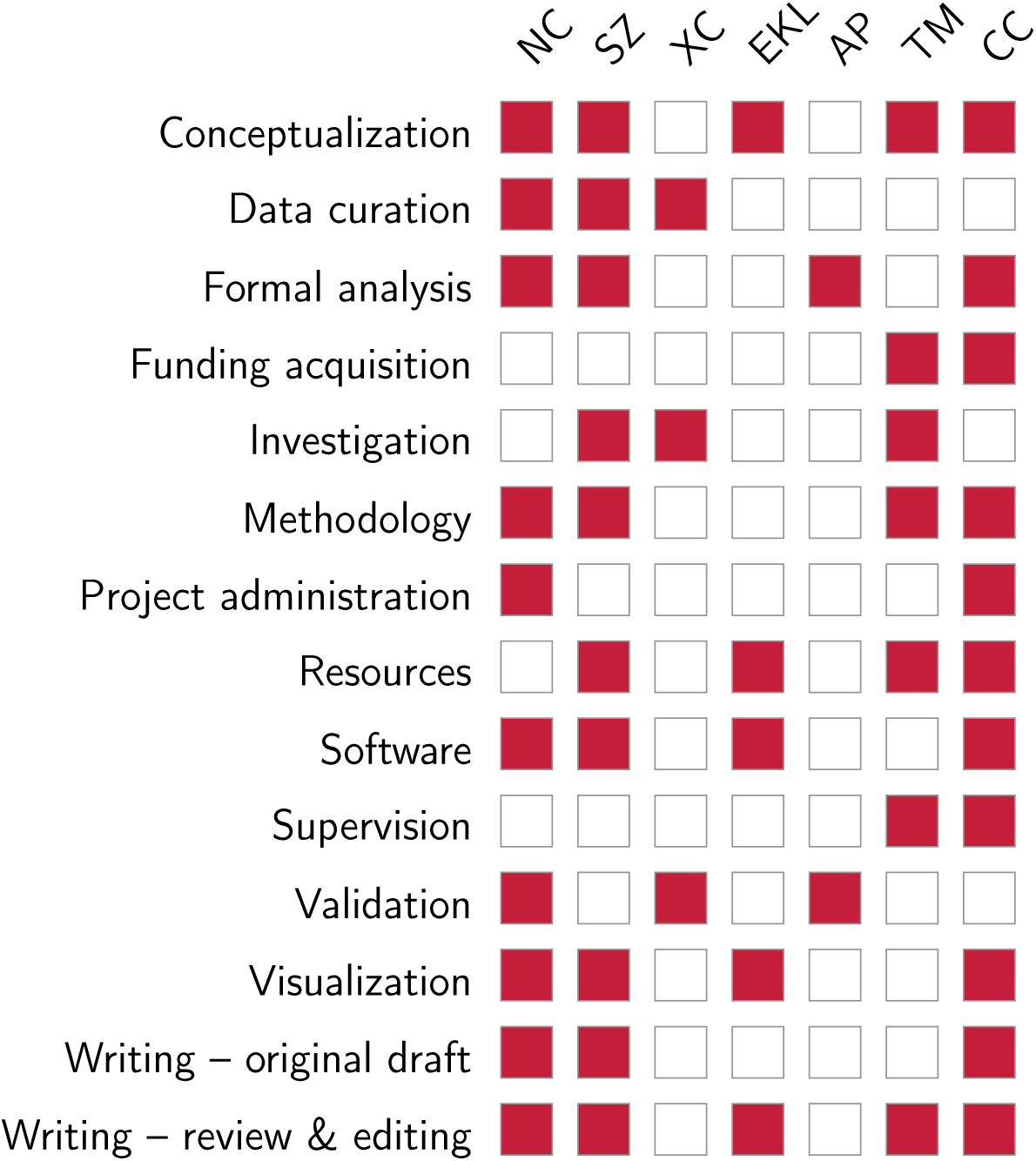

## 5.4. Data and Materials Availability

All data are available in the main text or the supplementary materials. The original session data is available online at Trepka E, Zhu S, Xia R, Chen X, Moore T (2022) Dryad Digital Repository Functional Interac-tions Among Neurons within Single Columns of Macaque V1 (https://doi.org/10.5061/dryad.x3ffbg7p2). All of the code used to analyze the data has been deposited to GitHub (https://github.com/carrn/V1_waveMAP) and is currently freely accessible online.

## Supplemental materials

**Figure S1:**
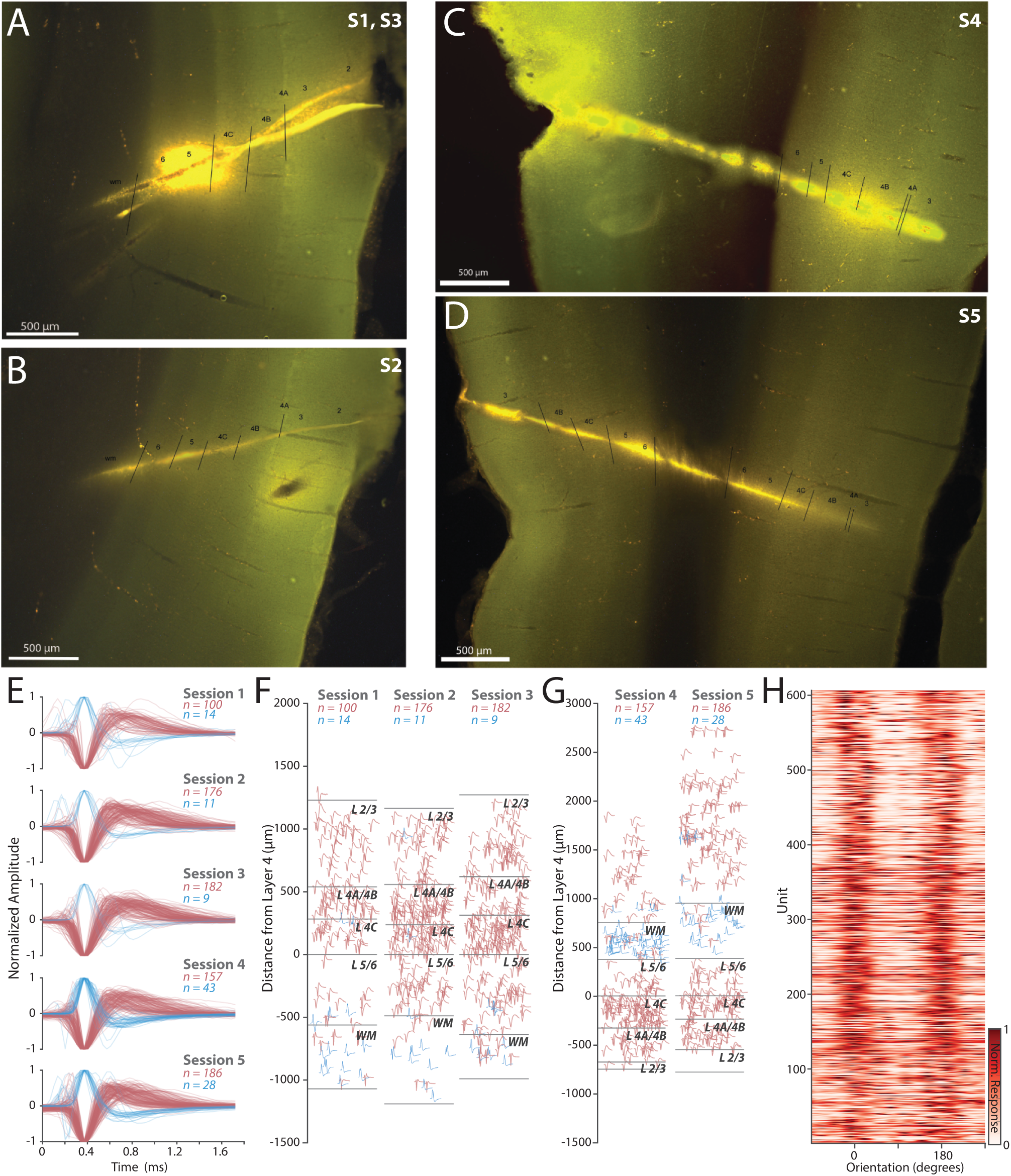
Histological verification of penetrations and neuronal waveforms from each session. (**A-D**) *Left*, Dye tracks of monkey 1 (M1) for sessions 1 and 3 (A), session 2 (B). *Right*, Dye tracks of monkey 2 (M2) for session 4 (C) and session 5 (D), all with delineations of laminar boundaries. (**E**) Curated waveforms for each of the five sessions. (**F-G**) Laminar organization of waveforms in sessions 1-3 (F, M1), and waveforms in sessions 4-5 (G, M2) (**H**) The normalized response tuning aligned to overall preferred orientation for 606 visually active neurons, sorted top to bottom by depth from superficial to deep.

**Figure S2:**
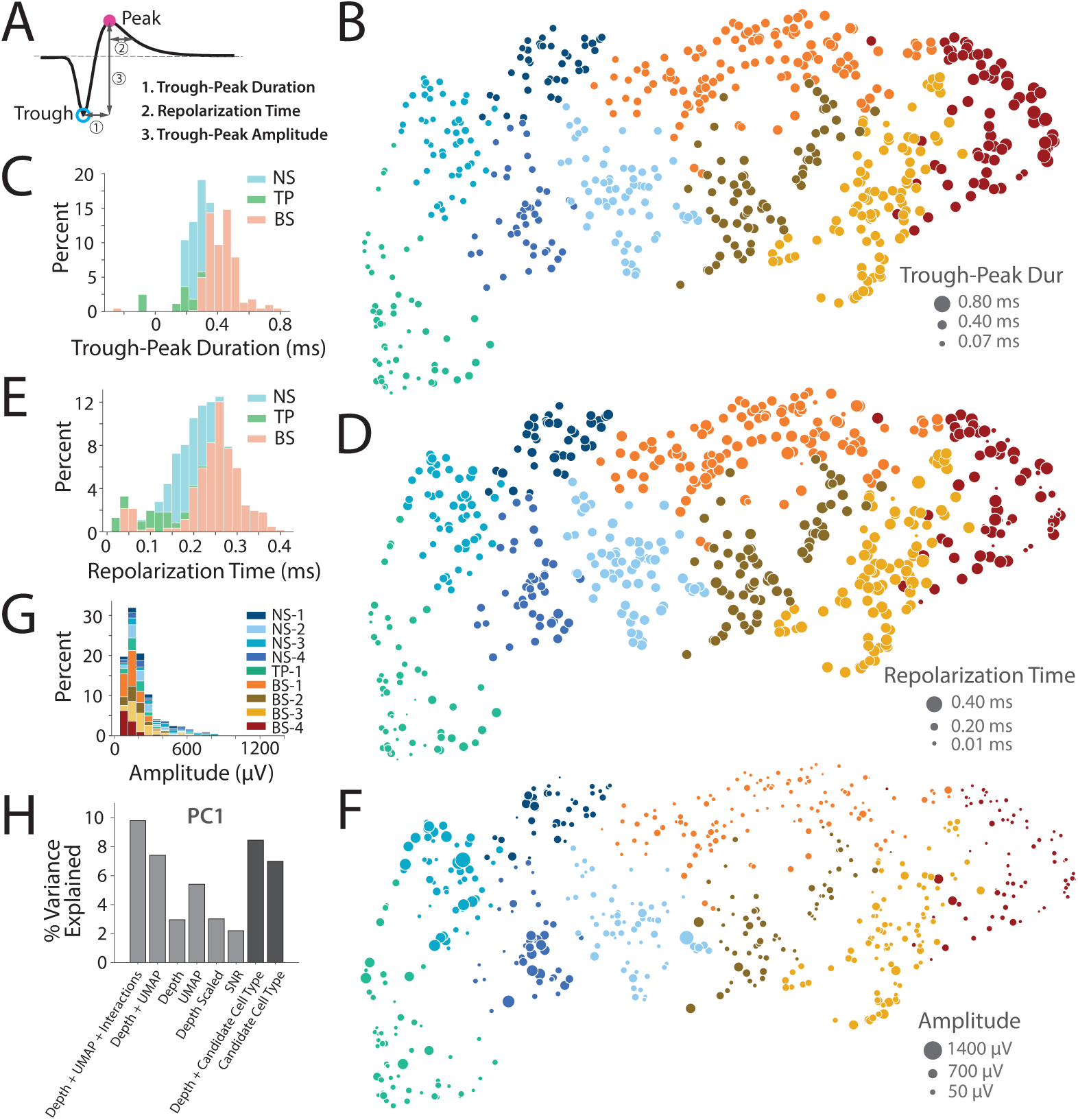
WaveMAP clusters lawfully reflect classic features. (**A**) Classic features of the extracellular waveform include peak (in pink solid circle), trough (in blue open circle), trough-peak duration (T-P Dur), repolarization time from 1/2 peak (Rep Time), and normalized trough-peak amplitude (T-P Amp). (**B**) UMAP X and UMAP Y plot with unit markers sized by trough-peak duration. UMAP X is strongly correlated with T-P Dur. (**C**) Histogram of all Trough-Peak Durations showing distributions of general categories (Tri-phasic, Narrow, and Broad). Note the distribution is unimodal and not easily separated into clusters. (**D**) UMAP X and UMAP Y plot with unit markers sized by 1/2 peak repolarization time. Again note strong correlation with repolarization time. (**E**) Histogram of all 1/2 peak repolarization times showing distributions of general categories (Tri-phasic, Narrow, and Broad). Note again the unimodal distribution of these features. (**F**) UMAP X and UMAP Y plot with unit markers sized by trough-peak amplitude, a variable not included in the UMAP analysis. The results suggest that some clusters have larger amplitudes than others. (**G**) Histogram of all amplitudes showing distributions for all clusters. (**H**) We performed PCA on the functional properties, orientation index, orientation circular variance, direction index, direction circular variance, orientation tuning bandwidth, and simple complex, of all visually responsive neurons to summarize the visual functional response of the neurons. We examined if this functional visual response could be explained by various predictors of cell type using a multiple linear regression. The predictors we used are UMAP coordinates, Depth, and various interaction terms between UMAP X coordinate and depth, UMAP Y coordinate and depth, and UMAP X, Y, and depth. As a control, we measured the amount of variance explained by signal-to-noise ratio to ensure our results were not a trivial artifact of spike sorting. Overall, both UMAP X and Y, Depth and candidate cell type predicted almost 10% of the variance in visual functional responses. Notably, the candidate cell type label and depth explained comparable amounts of variance as the depth and the UMAP coordinates.

**Figure S3:**
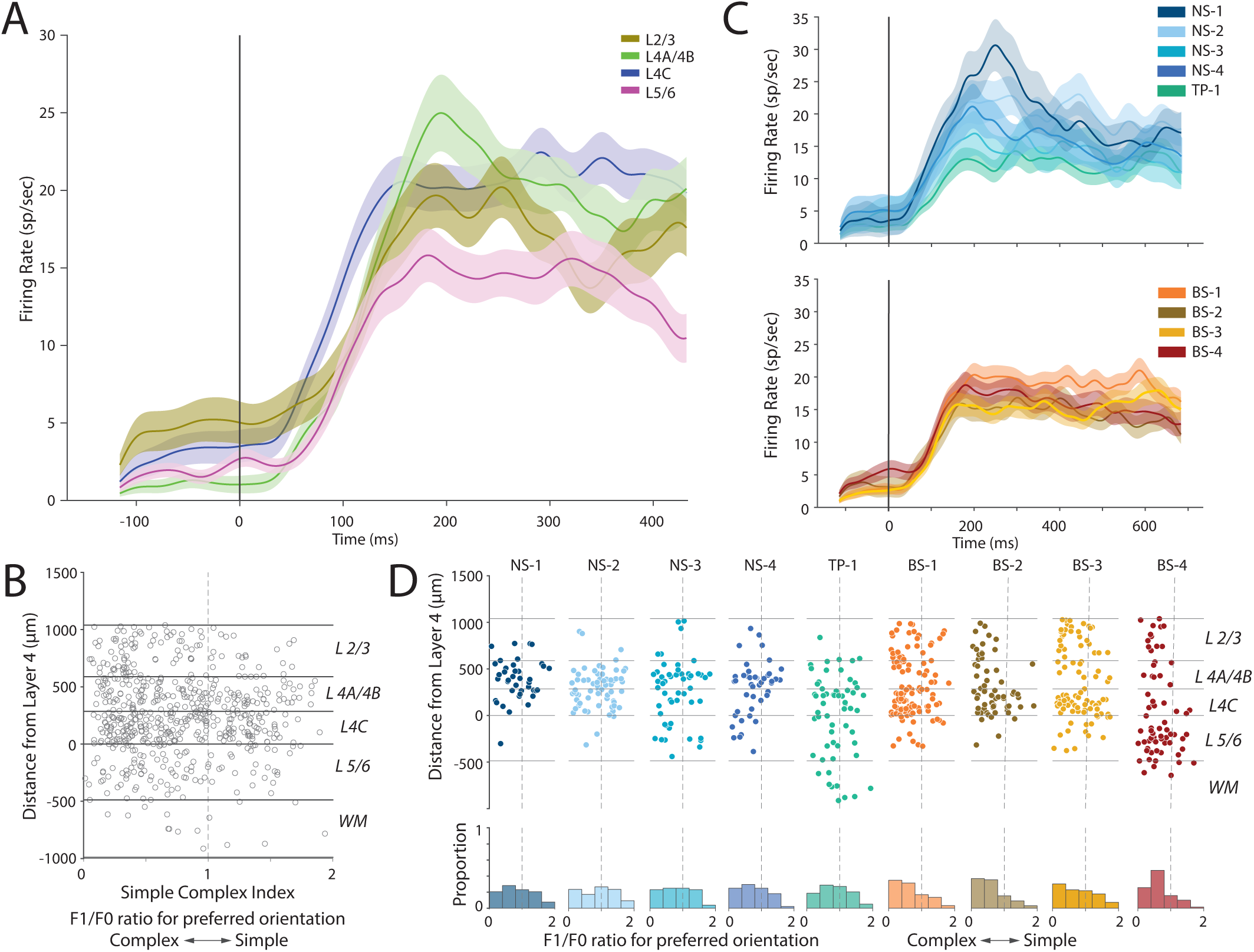
Neuropixels recordings in V1 are consistent with classical studies. (**A**) Average PSTH by layer. The black line indicates the initiation of the visual stimulus. (**B**) Scatter plot of modularity index (simple-complex index) as a function of scaled laminar depth of all units. Orientation selectivity is found in all layers but strongest in layer 4 consistent with classical reports (21). (**C**) Average PSTH by cluster. The black line indicates the initiation of the visual stimulus. *Top*, Mean PSTHs of “narrow” clusters. *Bottom*, Mean PSTHs of “broad” clusters. (**D**) There showed no strong indications that simple and complex cells mapped onto any one candidate cell type, although the broad-spiking clusters generally trended towards more complex responses. *Top*, Scatter plot of modularity index (simple-complex index) and scaled laminar depth separated by cluster. *Bottom*, Panel below the scatter plots shows histograms of the simple-complex index. The vertical dashed center lines show the boundary between complex (left of center) and simple (right of center).

**Figure S4:**
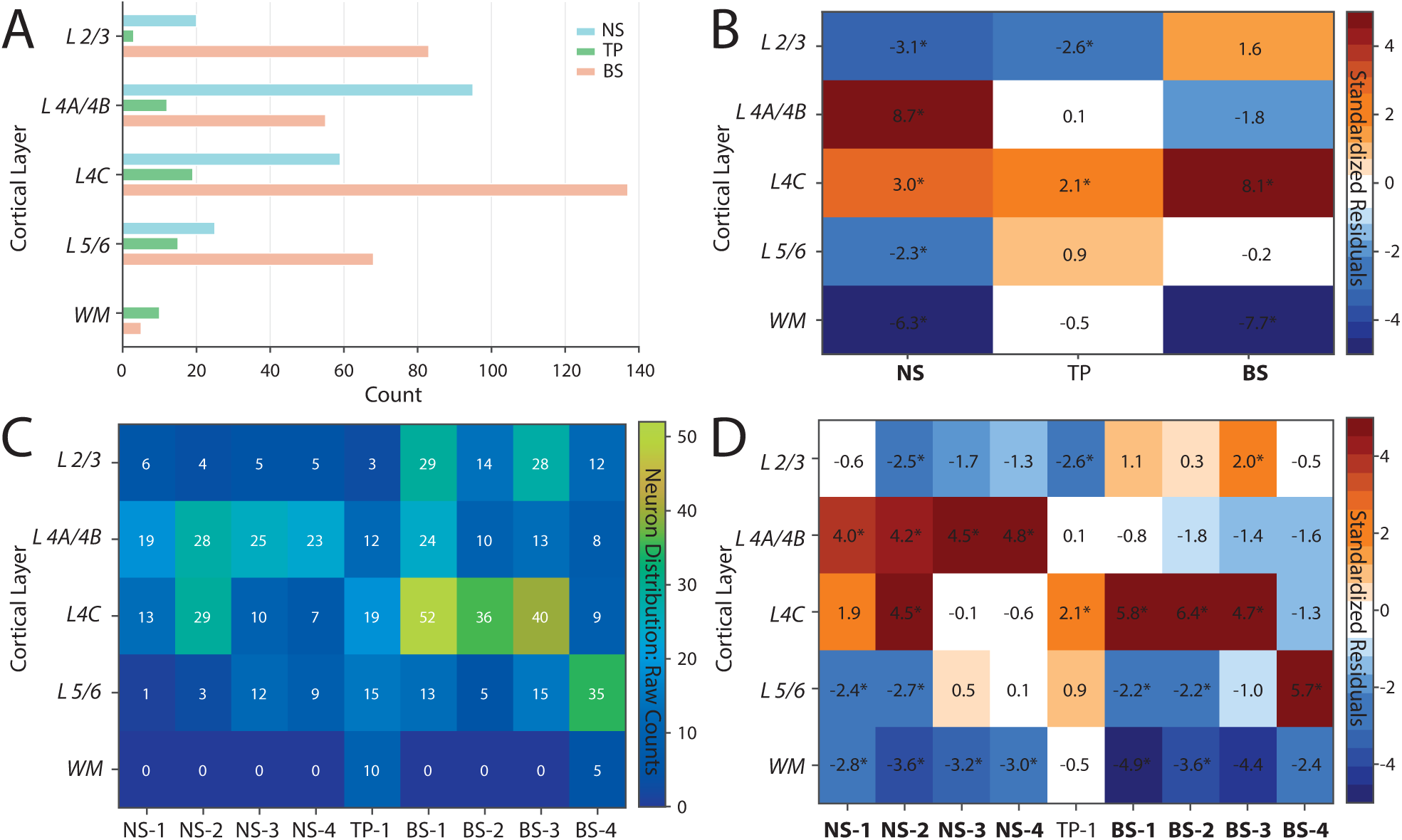
Narrow-spiking groups are concentrated in Layer 4A/4B/4C. (**A**) Histogram showing neuron count per layer separated by broad categories, narrow-spiking (NS), tri-phasic (TP), and broad-spiking (BS). (**B**) Under the null hypothesis that each layer (df = 4) should have the same proportion of neurons across the three broad categories (NS, TP, and BS), we performed a chi-square analysis to compare the observed distributions. Bold text indicates the category showed significance at p < 0.00001, otherwise indicates significance at p < 0.05. The heatmap shows standardized residuals, which are the difference between the observed and expected, divided by the square root of the expected count. * indicates a value ± 2.0 which suggests significance at p < 0.05. A positive value indicates a greater number observed than expected, and a negative indicates a lower number observed than expected. We see that the NS category is overrepresented in Layer 4A/4B and 4C. (**C**) Raw counts of neuron distribution by layer and cluster. (**D**) Under the null hypothesis that each layer (df = 4) should have the same proportion of neurons across each cluster, we performed a chi-square analysis to compare the observed distributions. Bold text indicates the cluster showed significance at p < 0.00001, otherwise indicates significance at p < 0.05. The heatmap shows standardized residuals, which are the difference between the observed and expected, divided by the square root of the expected count. * indicates a value ± 2.0 which suggests significance at p < 0.05. A positive value indicates a greater number observed than expected, and a negative indicates a lower number observed than expected. We see that all NS subtypes are overrepresented in Layers 4A/4B and NS-1 and NS-2 are overrepresented in Layer 4C.

**Figure S5:**
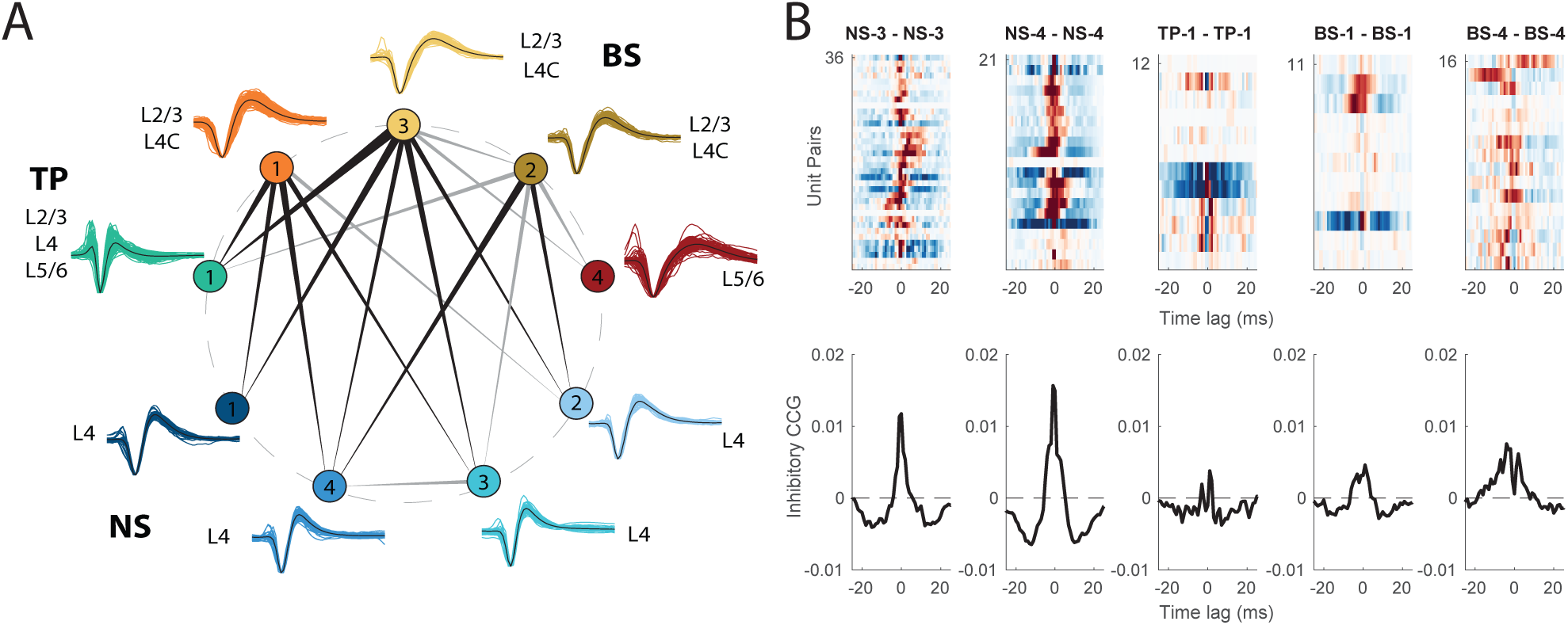
Excitatory and inhibitory cross-correlation analysis shows ordered connectivity between clusters. (**A**) Summary diagram of excitatory CCGs between clusters. Line thickness indicates the directionality of the CCG, from leading (thin line) cluster to lagging (thick line) cluster, with line width weighted by median lead-lag index. Black colored lines have p < 0.001, grey colored lines have p < 0.01. Overall, narrow-spiking neurons that are strongly localized to L4 and L5/6 lead broad-spiking neurons in L2/3 and L4, perhaps due to strong inputs into layer 4 neurons. (**B**) *Top*, Inhibitory CCGs between pairs of cells within each cluster. Only clusters which contained more than 10 pairs of cells are included. NS-3 and NS-4 show more inhibitory connections within their pairs than TP-1, BS-1, and BS-4. *Bottom*, Mean inhibitory autocorrelogram trace per each cluster.

**Figure S6:**
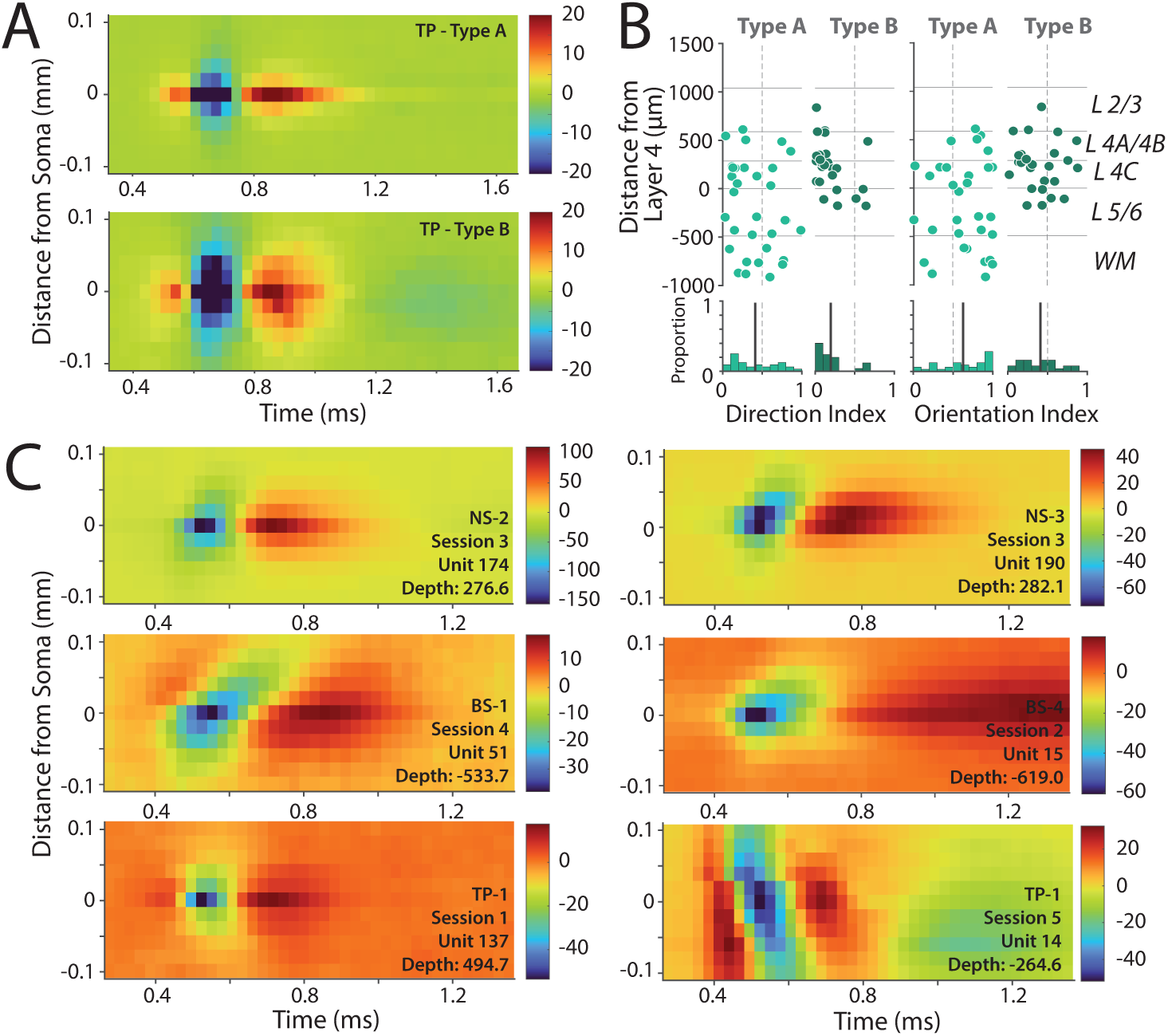
Spatial profiles show more hetereogeneity than cluster features typically capture. (**A**) Average multi-channel waveforms of tri-phasic cluster Type A (n = 32), and Type B (n = 25). Type A and Type B were manually classified on the criteria of spread (number of channels on which an extracellular action potential waveform continuum was observed). Of the 59 TP-1 Waveforms that were visually responsive, 2 were identified as artifactual. (**B**) *Top*, Laminar distribution of TP-1 Type A and Type B. The x-axis shows (from left to right) direction index and orientation index. A Wilcoxon rank sum test showed that the Type B population has significantly less direction selectivity (p < 0.01) and orientation selectivity (p < 0.05) than Type A. *Bottom*, Panel below the scatter plots shows histograms of the direction and orientation selectivity index. The bold line indicates the median orientation and direction index for each cluster. The vertical dashed center lines show the boundary between not selective (left of center) and selective (right of center). (**C**) Example multichannel waveforms from various clusters and sessions. *Upper Left*, “Narrow” Cluster 2 waveform. *Lower Left*, “Narrow” Cluster 6 waveform showing backpropagation. *Upper Right*, Tri-phasic Cluster 7 waveform. *Lower Right*, “Broad” waveform showing backpropagation.

**Figure S7:**
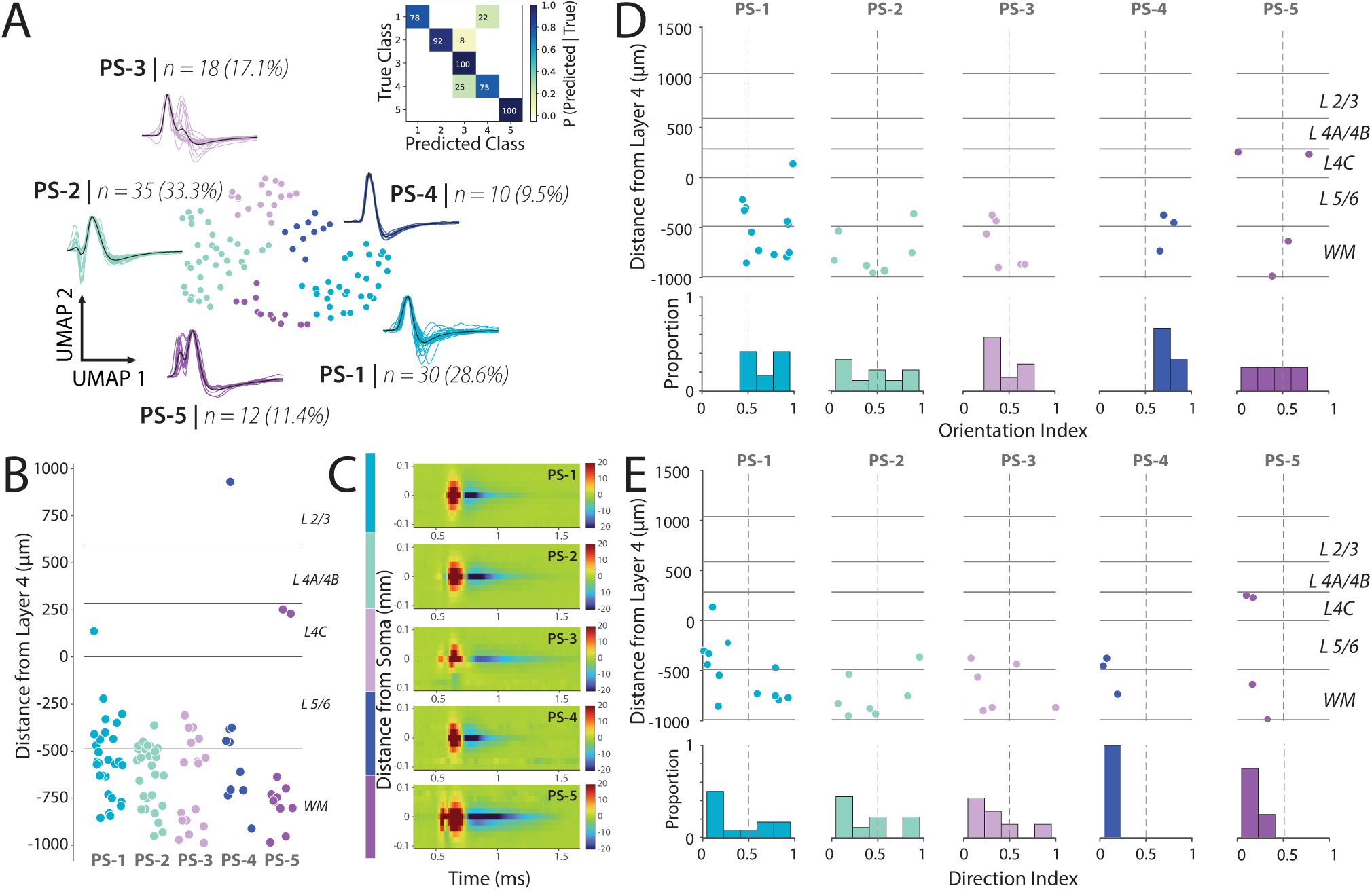
Positive waveform clusters are largely in white matter and show a diversity of visual responses. (**A**) Scatter plot of WaveMAP clustering on 104 positive spiking waveforms (N_neighbors = 20; MIN_DIST = 0.2; RESO-LUTION = 1.0) *Inset*, Confusion Matrix showing gradient boosted decision tree classifier with five-fold cross-validation. The main diagonal shows accuracy of waveform classification for each cluster, and off diagonals show misclassification percentages. (**B**) Laminar distribution of 5 positive spiking (PS) clusters across average laminar boundaries. Points are randomly jittered on the x-axis. (**C**) Average multichannel extracellular waveforms per cluster. Cluster color is shown on the left, and the colorbar shows the scale standardized for all clusters to map linearly between –20 and 20 *µ*v. (**D**) Of the 105 positive spiking units, 32 were responsive to visual stimuli. *Top*, Scatter plot of orientation index and scaled laminar depth separated by cluster for positive spiking units. *Bottom*, Panel below the scatter plots shows histograms of the orientation index. The vertical dashed center lines show the boundary between not selective (left of center) and selective (right of center). While the numbers are too small to strong statistical conclusions, clusters PS-1 and PS-4 have strong orientation selectivity but low direction selectivity similar to TP-1 type B neurons (Fig. S6B). (**E**) *Top*, Scatter plot of direction index and scaled laminar depth separated by cluster for positive spiking units. *Bottom*, Panel below the scatter plots shows histograms of the direction index. The vertical dashed center lines show the boundary between not selective (left of center) and selective (right of center). In general, clusters PS1-5 show modest direction selectivity.

**Table S1:**
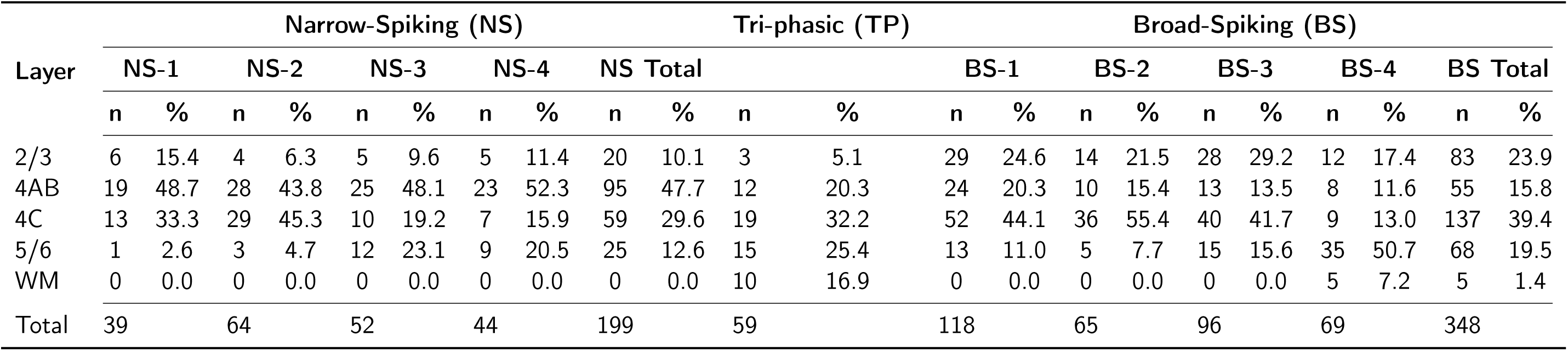
Distribution of Neuronal Clusters Across Cortical Layers.

